# Single cell transcriptomics reveals involution mimicry during the specification of the basal breast cancer subtype

**DOI:** 10.1101/624890

**Authors:** Fatima Valdes-Mora, Robert Salomon, Brian Gloss, Andrew MK. Law, Lesley Castillo, Kendelle J. Murphy, Jeron Venhuizen, Astrid Magenau, Michael Papanicolau, Laura Rodriguez de la Fuente, Daniel L. Roden, Yolanda Colino-Sanguino, Zoya Kikhtyak, Nona Farbehi, James RW. Conway, Samantha R. Oakes, Neblina Sikta, Seán I. O’Donoghue, Thomas R Cox, Paul Timpson, Christopher J. Ormandy, David Gallego-Ortega

## Abstract

Both luminal and basal breast cancer subtypes originate in the mammary luminal progenitor cell compartment. Basal breast cancer is associated with younger age, early relapse, and high mortality rate. Here we used unbiased droplet-based single-cell RNAseq to elucidate the cellular basis of tumour progression during the specification of the basal breast cancer subtype from the luminal progenitor population. Basal–like cancer cells resembled the alveolar lineage that is specified upon pregnancy and showed molecular features indicative of an interaction with the tumour microenvironment (TME) including epithelial-to-mesenchymal transition (EMT), hypoxia, lactation and involution. Involution signatures in luminal breast cancer tumours with alveolar lineage features were associated with worse prognosis and features of basal breast cancer. Our high-resolution molecular characterisation of the tumour ecosystem also revealed a highly interactive cell-cell network reminiscent of an involution process. This involution mimicry involves malignant education of cancer-associated fibroblasts and myeloid cell recruitment to support tissue remodelling and sustained inflammation. Our study shows how luminal breast cancer acquires an aberrant post-lactation developmental program that involves both cancer cells and cells from the TME, to shift molecular subtype and promote tumour progression, with potential to explain the increased risk and poor prognosis of breast cancer associated to childbirth.

## Background

The mammary gland is a unique organ that undergoes a series of developmental processes that mostly occur postnatally, with profound tissue morphogenesis during puberty and pregnancy[1]. During pregnancy, alveolar milk-secretory epithelial cells differentiate from the CD61+ luminal progenitor population (also EpCAM^high^/Sca1^+^/CD49b^+^)[2, 3] driven by Prolactin- and Progesterone-induced up-regulation of Elf5, the master transcriptional regulator of the alveolar cell fate[4-6]. Transcriptional profiling at single-cell resolution has recently shown however, that this differentiation process is more complex with multiple subtypes and states involved in the differentiation of the alveoli during pregnancy[7].

Tissue morphogenesis associated with pregnancy and lactation concludes with the developmental process of mammary involution, a process whereby the lactating mammary gland returns to a quasi-nulliparous state[8, 9]. Involution is a highly complex multicellular process where milk stasis triggers alveolar cell death and apoptosis[10]; neutrophils and macrophages subsequently engulf the death cell bodies of the collapsing alveoli and regulate inflammation by suppressing the adaptive immune response; finally, fibroblasts are activated to fill the areas left by the dying epithelium in a pseudo-wound healing process[11] and an intense adipogenesis program is activated. This exquisitely regulated morphogenesis of the mammary gland requires a high degree of cell-to-cell communication within the mammary epithelium and throughout different cell types from the tissue microenvironment.

The contribution of normal process during mammary gland development to breast cancer has been under intense research in the last few years. Some mammary epithelial cell lineages have been proposed as the cell of origin of the different breast cancer subtypes[12, 13], including those with specific genetic aberrations[13, 14]. The association of developmental cues with breast cancer is not restricted to the epithelial compartment, as tissue morphogenesis encompasses dramatic changes within the epithelial ductal tree as well as its surrounding stroma. Thus, these events of tissue remodelling are often hijacked in breast cancer as drivers of tumour progression[15]. This might explain the transient higher risk of developing breast cancer during the 10 years after childbirth. Importantly, this pregnancy-associated breast cancer (PABC), correlates with metastatic disease and poor prognosis and is generally associated with younger women (reviewed in[16]). Despite the effects of the hormonal load associated with pregnancy, there is increasing evidence pointing to a direct effect of the TME and the events of tissue remodelling that occur in the mammary gland during pregnancy, and in particular those associated to involution, as the driving force behind the higher mortality associated to PABC[11, 16, 17].

We have previously shown that pregnancy-associated transcriptional networks driven by the transcription factor Elf5 specify the basal subtype of breast cancer, promoting a lethal phenotype, characterised by endocrine insensitivity and resistance to therapy[18] and the promotion of metastatic disease through a profound alteration of the TME[19]. Here, we use spatially and time controlled induction of Elf5 within the mammary epithelium in the context of the luminal MMTV-PyMT mouse mammary cancer to study tumour progression mechanisms associated to the specification of the basal breast cancer subtype at single-cell resolution. This study provides an unprecedented characterisation of the intense TME remodelling acquired by the aberrant activation of pregnancy-related developmental pathways during breast carcinogenesis. Our analysis revealed the cellular diversity of breast tumours, characterising cell subclasses that resemble the mammary epithelial lineages and identifying strong plasticity within the luminal progenitor compartment, the origin of luminal and basal cancers. Besides the contribution of cell-intrinsic cues on the alveolar cell differentiation to breast cancer progression [18, 20], we exposed the cell-extrinsic effects on the TME[19], such as EMT, collagen deposition, inflammation, vascular leakiness and hypoxia. Using our high-resolution molecular landscape of breast tumours, we mapped cell-to-cell interactions within the TME, revealing a multicellular mechanism of tumour progression that encompasses immune suppression and tissue remodelling, two central processes involved in the promotion of metastatic disease[19]. We demonstrate that this mechanism of tumour progression is subsequent to the alveolar cancer cell lineage specification and it is linked to a mimicry involution process. Finally, we provide evidence at the single-cell resolution of the molecular pathways that alveolar cells elicit to recruit Myeloid-Derived Suppressor Cells (MDSCs) and activate specific cancer-associated fibroblasts (CAFs) subpopulations; two central players in this process of involution mimicry.

## Results

### Unbiased massively-parallel scRNAseq captures cell heterogeneity of MMTV-PyMT mammary tumours

In this study we used two mouse models, the well-established MMTV-PYMT mouse mammary tumour model^25-27^ (PyMT/WT) and a mammary restricted (MMTV) doxycycline-inducible (rtTA) Elf5 expression model[4, 19] crossed with the PyMT/WT model[19, 21-23] (PyMT/Elf5) (SuppFig.1A). Consistent with previous reports, the specific induction of Elf5 in the mammary gland epithelium results in a significant increase of lung metastases and a dramatic leaky vasculature phenotype in the PyMT model (SuppFig.1B-D and Supp Videos 1 and 2)^18^. A total of fifteen mouse mammary tumours at metastatic stage (14 weeks old FBVn background) from the MMTV-PyMT/WT and /ELF5 model were harvested and analysed using Drop-seq[24]. After quality control an filtering, a total of 15,702 cells were analysed (see material and methods and SuppFig. 2A-B). Cells from the biological replicates contributed to all cell clusters, indicating consistent cell sampling between all replicates (supplementary information and SuppFig. 2C) and the expression of Elf5 was higher in PyMT/Elf5 tumours SuppFig. 2D). We then annotated the main cellular compartments in the tumours using cell signatures defined by Aran and colleagues[25] (Fig. 1A). The majority of cells were defined as epithelial cells (63.16%); tumour associated stroma represented 20.41% of the cells identified; and the tumour-infiltrated immune cell compartment accounted for 16.44%. This classification was consistent with the cell surface marker analysis (FACS) to define epithelial cells, leukocytes and stroma cells (Fig. 1B). A tSNE visualisation of the three cellular compartments and their associated top expressed markers is shown in Figure 1C; and we used the RNA expression of canonical markers for each compartment to verify their identity (SuppFig. 3A). Consistent with the role of Elf5 as a master regulator of mammary cell specification [4, 5], we found a striking shift in the main epithelial cluster associated to the sample genotype (Fig. 1D). This result clearly shows that induction of ELF5 expression in PyMT tumours generate a profound transcriptional redefinition of the epithelial cell compartment, this effect was not observed in the other cellular compartments.

**Figure 1.**
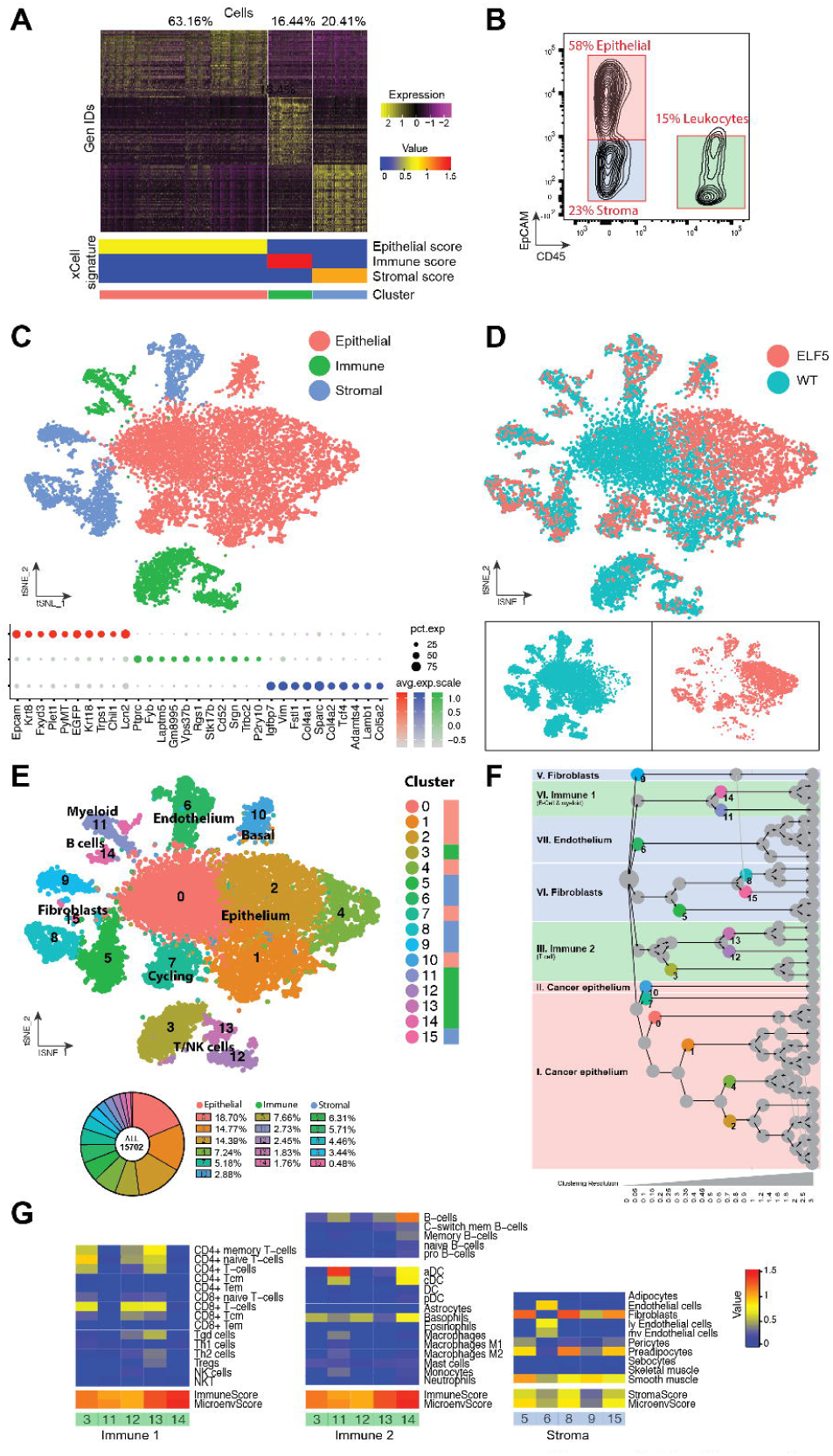
High-resolution cell composition of MMTV-PyMT tumours. **A)** Heatmap showing differential expression of the top expressed genes contributing to the epithelial (red), stromal (blue) and immune (green) clusters and their percentage. The bottom panel shows the enrichment value for each of the signatures of the xCell algorithm for each cluster is shown. **B)** Representative contour plot of the cell composition of a MMTV-PyMT tumor analysed by flow cytometry defined by EpCAM antibodies (epithelial cells), CD45 (leukocytes) and double negative cells (stroma). **C)** tSNE visualisation shows the coordinates of each analysed cell after dimensional reduction coloured by its main cell lineage. The dot plot shows the top differential markers form each of the main cell lineages and their level of expression. **D)** tSNE plot showing the distribution of cells per genotype. **E**) tSNE plot showing cell clusters defined by a k-means based clustering algorithm, and their relative frequency (bottom). **F)** Cluster tree modelling the phylogenic relationship of the different clusters in each of the main cell types compartments at different clustering resolutions. Dashed red line shows the resolution chosen. Coloured circles in the cluster tree represent the origin of the clusters represented in the tSNE plot shown in panel E. **G)** Cell identification using score values for each of the metasignatures of the xCell algorithm in the immune and stromal compartment divided by cluster.

We then used differential gene expression analysis and k-means clustering to identify cell diversity within the major cell compartments, classifying the tumour cells into 16 different cell classes (Fig. 1E). A cluster tree representation[26] of the relationship among the different clusters revealed two main families in the immune compartment, three in the stromal compartment and two in the cancer compartment; all of those defined very early in the phylogeny (Fig. 1F). The top differential marker genes that define stromal and immune cell compartments is shown in SuppFig. 3B. Such cell diversity within the major compartments was also validated using xCell algorithm [25] (Fig. 1G). The xCell signatures clearly identified endothelial cells and various classes of fibroblasts in the stromal compartment; a clear segregation between T-cells and all the other immune species such as B-cells and the diverse array of myeloid cells.

In conclusion unbiased Drop-seq identifies cell heterogeneity in all three main compartments of epithelial, stroma and infiltrated immune cells from PyMT tumours.

### PyMT cancer cells are diversely organised in a structure that resembles the mammary gland epithelial hierarchy

In our Drop-seq analysis, the major change by ELF5 induction was confined to the epithelial cell compartment (Fig. 1D). Thus, we performed an independent k-means clustering on the epithelial subset cohort of cells (6,475 PyMT/WT and 3,233 PyMT/Elf5 cells) at an increased resolution. Epithelial tumour cells were distributed in 11 clusters visualised using tSNE representation (Fig. 2A). Figure 2B shows the strong genotype-driven cell enrichment in the epithelial compartment, which was not due to batch effects (SuppFig. 4A and B). The tSNE plot from Figure 2B shows that clusters 1 and 6 (C1 and C6) were formed mostly in response to forced ELF5 expression (>2 Fold of cells belong to PyMT/ELF5 mice), while C0, C7, C9, C4 and C3 were formed mostly by WT cells (>2 Fold of cells belong to PyMT/WT mice) (Fig. 2C). C2, C5, C8 and C10 were present in both WT and forced ELF5 expressing tumours (<2 Fold of cells from any of the genotypes). This distribution is consistent with the expression of ELF5 in each of these clusters (SuppFig. 4C). Concomitantly with the scRNAseq data, FACS analysis of the cancer epithelial cells revealed that Elf5 forced differentiation of the luminal progenitors, defined by a decrease in the proportion of Sca1^low^/CD49b^+^ progenitor cells[2] within the luminal (EpCam+/CD49f+) population (Fig 2D).

**Figure 2.**
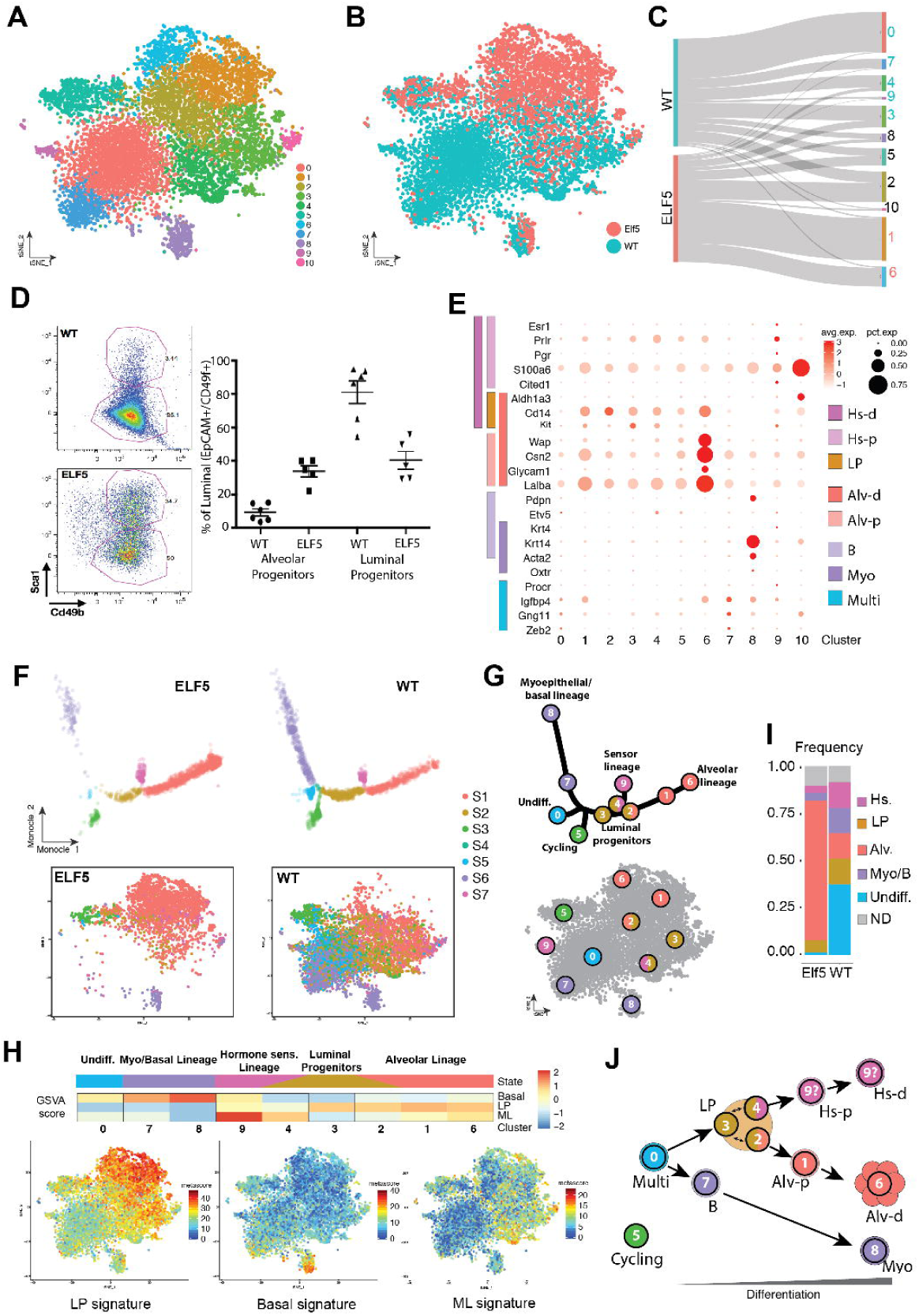
Cancer epithelial cell diversity of PyMT tumours are structured in lineages that resemble the mammary gland epithelial hierarchy. **A)** tSNE visualisation of the epithelial cell groups defined by k-means clustering analysis. **B)** Distribution of cells by genotype in the defined tSNE dimensions. **C)** Sankey diagram showing the contribution of each of the genotypes to each cell clusters. Cluster numbers are coloured by the dominant genotype (>2-fold cell content of one genotype), Elf5 (red), WT (green). **D)** Scatter plot showing flow cytometry data to define the % alveolar versus luminal progenitors using canonical antibodies that define the epithelial mammary gland hierarchy (EpCAM, CD49f, Sca1 and CD49b), in PyMT tumours (WT n = 6 and Elf5 n = 5). Bottom panels are representative flow cytometry plots of one of the replicates for each genotype. **E)** Dot plot representing the expression level (red jet) and the number of expressing cells (dot size) of the transcriptional mammary gland epithelium markers in each PyMT epithelial cluster. These marker genes were grouped according to each mammary epithelial cell type as defined by Bach, et al.: Hormone sensing differentiated (Hs-d, dark pink), Hormone-sensing progenitor (Hs-p, light pink), Luminal progenitor (LP, orange), Alveolar differentiated (Alv-d, dark red), Alveolar progenitor (Alv-p, light red), Basal (B, light purple), Myoepithelial (Myo, dark purple), Undifferentiated (Multi, light blue). **F)** Top panels: trajectory analysis of the PyMT/WT and /ELF5 cancer epithelial cells along the pseudotime signatures that define the main lineages of the mammary gland epithelial hierarchy. Bottom panels: projection of the states defined by pseudotime analysis into tSNE clustering coordinates overall and per trajectory state. **G)** Schematic representation of the overlayed cell states (S) and cluster cell identities (C). **H)** Enrichment analysis (GSVA score) for the gene signatures that define the main mammary gland lineages: Basal, Luminal Progenitor (LP) and Mature Luminal (ML) for each of the clusters. Bottom tSNE representations show the expression of each of the gene signatures at single cell resolution. The top bar shows the assigned mammary epithelial cell type as per section C. **I)** Frequency of the different cell lineages in each genotype. **J)** Illustration of cell diversity of PyMT tumours based on the canonical structure of the mammary gland epithelial lineages.

We then annotated the cells from our MMTV-PyMT epithelial clusters with the normal cells from the mammary epithelium, using the defined canonical markers by Bach and colleagues [7] (Fig. 2E). C0 and C7 expressed genes consistent with Procr+ multipotent/stem cells. C9 cells expressed hormone sensing (Hs) lineage markers, C8 expressed markers for Basal (B)/ Myoepithelial (Myo) cells, whereas C1, C2, C3, C4, and C6 presented markers consistent with luminal cells, with C2, C3, and C4 presenting progenitor markers while cluster C1 and C6 presented markers consistent with alveolar differentiated cells. C5 and C10 were the most undefined clusters, showing mixed luminal and myoepithelial markers C5 corresponded with cycling cells (SuppFig. 4D) and C10 showed very low levels of the PyMT oncogene (SuppFig. 5A), suggesting an untransformed origin of normal cells trapped within the tumour mass. A complete functional annotation of each epithelial cluster using Gene Set Variation Analysis (GSVA) can be found in (Supp. Table 1, SuppFig. 5 and Supplementary Information)

### Dynamic states of the malignant lineages of PyMT tumours: implications for the cell of origin of cancer

To further investigate the differentiation states present within the luminal lineage in the PyMT tumours, we performed pseudotime analysis using Monocle. We used the mammary gland hierarchy gene signature [27] to build the pseudotime structure of the PyMT cancer epithelial cells identifying seven discrete states (Fig. 2F, upper panels). Consistent with previous results, Elf5 had a profound impact upon the structure of the epithelial cell diversity biasing the composition of the cell lineages towards one of the end fate axis (right hand side) of the pseudotime plot dominated by state 1 (S1) at the expense of a depletion of states S2, S4, S5 and S6. We then visualised the Monocle defined cell states using the coordinates from the tSNE and viceversa, which allowed us to overlap the cell cluster identity (C) with the pseudotiming states (S) (Fig. 2F bottom panels, Fig. 2G).

The alveolar lineage state (red, S1) clearly corresponded to 2 distinct cell identities (cell clusters C1 and C6); the sensor lineage state (pink, S7) was composed of C9, and the myoepithelial/basal lineage state (purple, S6) was formed by C7 and C8. An undifferentiated state with multipotent/stem characteristics (blue, S5), which correspond to C0, sat in a position equidistant from the myoepithelial and the luminal lineages (alveolar and sensor). Interestingly, and consistently with the literature where stem cell properties are confounded by myoepithelial properties [28], C7 was classified by Monocle as part of the basal/myoepithelial lineage, however, this cluster presents strong multipotent characteristics similar to those present in C0. The most difficult compartment to define was the luminal progenitor state (brown, S2, represented by C2, C3 and C4) presumably due to their strong plasticity and subtle differential features.

Finally, to further gain insights in the definition of the epithelial clusters, we used GSVA [29] and metasignature visualisation to score each of the clusters according to the mammary gland epithelial signatures defined by Pal and colleagues. (Fig. 2H). Hierarchical clustering based on GSVA scores associated C2 with the alveolar lineage, suggesting its identity as a pre-alveolar committed luminal progenitor, or a cell type that sits in the hinge between the luminal progenitor and the alveolar lineages. Similarly, C4 was associated to the hormone-sensing lineage represented by C9, indicating a luminal progenitor committed towards this lineage. In this scenario, C3 remained undefined so we classified it as uncommitted luminal progenitor. tSNE visualisations of these metasignatures are shown in Fig 2H which further confirmed the cell distribution of the three main mammary epithelial lineages. The distribution of the different designated lineages in the two genotypes is shown in Fig. 2I, showing less heterogeneity in PyMT/ELF5 tumours induced by a strong enrichment of the alveolar lineage.

Altogether, the combination analysis of gene signatures, gene markers and pseudotiming enabled the precise annotation of PyMT cell clusters within the mammary hierarchy, identifying a large luminal lineage that retains most of the cell diversity and strong plasticity, a basal/myoepithelial compartment and a hormone-sensing lineage (Fig. 2J).

In conclusion, and consistent with previous transcriptomic classification using bulk RNAseq [13], we found that the cell composition of PyMT tumours is dominated by luminal progenitors. However scRNAseq reveals a more complicated structure as multiple lineages and their intermediate cell progenitors are found in these tumours. Elf5/PyMT tumours on the other hand, showed more homogenous tumours dominated by the alveolar lineage.

### Molecular mechanisms of cancer progression associated to cancer cells of Alveolar origin

We next sought to validate some of the functional findings found in cancer cells of alveolar origin (Clusters 1 and 6) characteristic of PyMT/ELF5 tumours, including the decrease of cell proliferation, lack of EMT [18-20] and increase of hypoxia (SuppTable 1, SuppFig. 5 and Supplementary Information). Figure 3A shows the distribution of cell cycle marker signatures [30] in the epithelial cell compartment. These signatures identified a specific group exclusively populated by cells in G2M or S phase of the cell cycle. PyMT/ELF5 tumours showed an enrichment in the G1 phase, this is consistent with our previous reports that ELF5 increased cell cycle time by delaying entry into S phase [18], and that ELF5 reduced cell proliferation in PyMT tumours [19].

**Figure 3.**
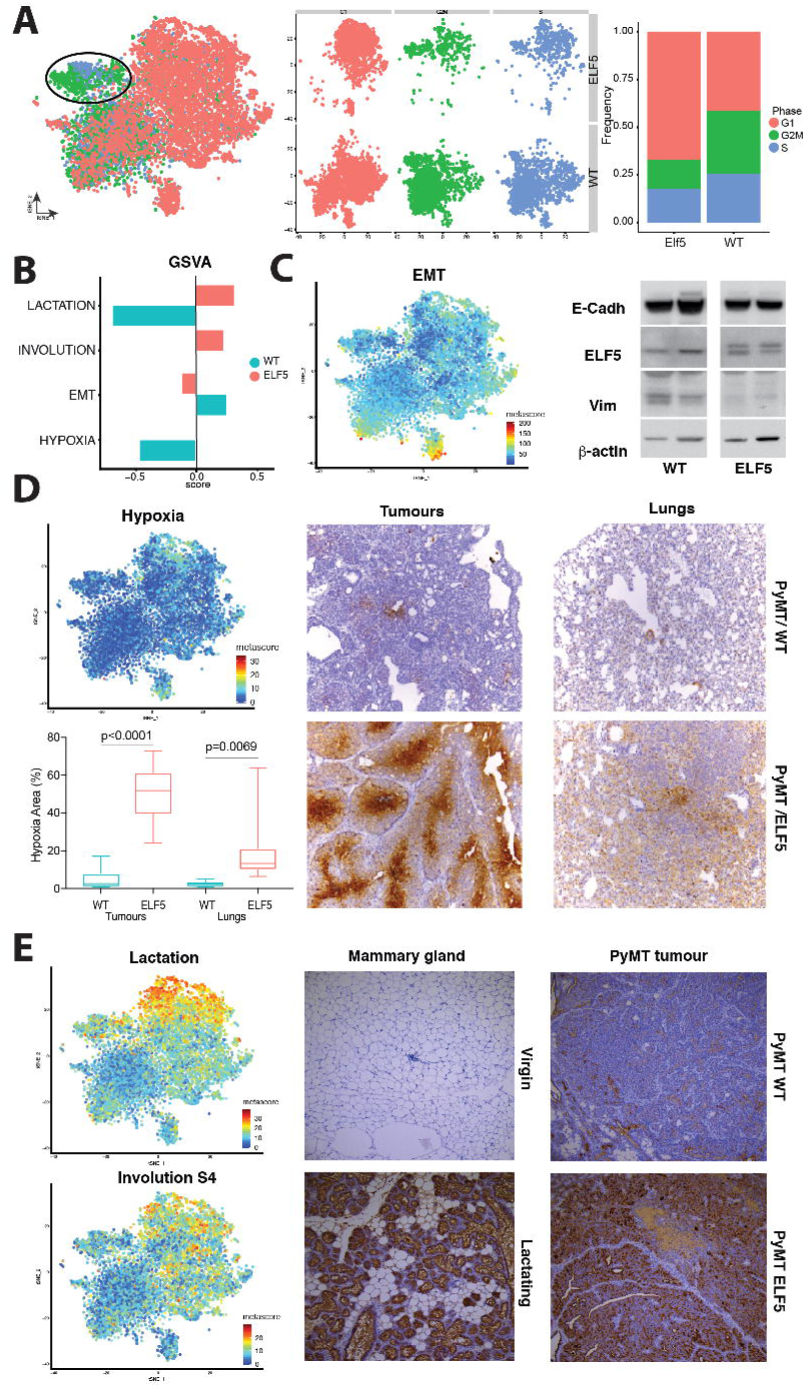
Functional validation of cancer related features associated to PyMT/Elf5 tumours. **A)** Cell cycle stages of the PyMT cancer cells as defined by gene expression signatures using tSNE coordinates and their deconvolution. Circled area shows the cycling cluster (C5 in Fig. 2) characterised by a total absence of G1 cells. The quantification of the proportion of cells in each stage grouped by genotype is shown in the bar chart. **B)** Enrichment GSVA analysis of gene expression metasignatures of cancer-related and Elf5-related hallmarks associated to PyMT/WT (green) and /Elf5 (red) tumours. **C)** tSNE representation of the EMT gene expression metasignature. Right panel shows a western blot of canonical EMT markers (E-Cadh, E-Cadherin and Vim, vimentin) on PyMT/WT or ELF5 full tumour lysates. Note: the two images correspond to the same western blot gel cropped to show the relevant samples. **D)** Hypoxia metasignature is shown in the tSNE plot, bottom panel shows a bar plot of the extension of the hypoxic areas in PyMT/WT (green) and /Elf5 (red) tissue sections (tumours and lung metastasis) stained using IHC based on hypoxyprobe binding, representative images are shown in the right panels. **E)** Lactation and Late Involution (stage 4) metasignatures is shown in the tSNE plots. Pictures show IHC with an anti-milk antibody in tissue sections from a lactating mammary gland at established lactation compared with a mammary gland from an aged-matched virgin mouse; and in PyMT/WT and /Elf5 tumours.

We then performed GSVA for specific signatures of known functions of ELF5, and functions suggested by the Hallmark analysis from SuppFig. 5 and SuppTable 1. This analysis revealed a strong enrichment of lactation, involution, and hypoxia pathways and a decrease in epithelial-to-mesenchymal transition (EMT) in ELF5 tumours (Fig. 3B), which it is a known characteristic of ELF5 [18-20]. Visualisation at the single cell resolution of the EMT gene signature confirms this result with a strong EMT association only with myoepithelial and undifferentiated cells (Fig. 3C). Western Blot on total cell lysates of PyMT/WT and /ELF5 tumours for canonical markers of EMT that have been previously identified to be regulated by ELF5 via repression of SNAI2 [20] confirmed that ELF5 tumours were depleted of such EMT markers at the protein level. Although E-Cadherin levels were not significantly different, ELF5 tumours with alveolar features show a clear reduction of Vimentin (Vim) indicating MET (Fig. 3C).

PyMT/ELF5 tumours presented extensive vascular leakiness (SuppFig. 1D), a fact that can lead to increased hypoxia [19]. Cells enriched for a hypoxic gene signature concentrated in myoepithelial and alveolar cells (Fig. 3D, SuppFig. 7 and SuppTable 1). We confirmed these findings using a hypoxyprobe to identify hypoxic regions in the tumours. Immunohistochemistry showed extensive hypoxic regions in PyMT/ELF5 tumours compared to their WT counterparts, indicating that cancer alveolar cells are strong drivers of events leading to hypoxia or these cells are able to survive better in hypoxic regions. This phenotype was strong enough to induce hypoxia in the metastases of a highly oxygenated tissue such as the lung (Fig. 3D).

As expected, the lactation signature was strongly enriched in cells of alveolar origin (Fig. 3E, top tSNE plot). Immunohistochemistry using anti-milk antibodies revealed a strong production of milk in PyMT/ELF5 tumours compared to PyMT/WT tumours (Fig. 3E, IHC pictures). In the normal mammary gland, milk stasis generates the signals that trigger mammary involution, and in this scenario, where milk cannot leave the tumour, we identified a late involution gene signature [31] associated with the luminal lineage in PyMT tumours, specifically enriched in ELF5 alveolar cells (Fig. 3E, bottom tSNE plot).

Taken together, single-cell transcriptomics enables the identification of biological properties of mammary tumours and, in particular, the role of involution mimicry as a plausible driver of sustained inflammation associated with the Elf5-alveolar subtype in luminal breast cancer.

### Characterisation of cancer-associated fibroblasts in PyMT tumours

Involution is a multistep and multicellular process that involves alveolar cell death and tissue remodelling, the latter is orchestrated by stromal and immune cells [32]. Among these cell types, fibroblasts have a critical role during the extracellular matrix remodelling and immune suppression steps [33]. Thus, we further explored the role of cancer-associated fibroblasts (CAFs) in aberrant involution in our model as a potential key event that could fuel Elf5-driven metastatic mammary tumours.

Unsupervised clustering on the fibroblast subset (2,255 pooled cells from PyMT/WT and PyMT/ELF5 tumours) revealed three major fibroblast clusters (Fig. 4A, SuppFig. 6A-B). The overlay of the cell cycle signature in the tSNE showed that a cell subgroup of fibroblast cluster 2 corresponded to dividing cells (SuppFig. 6C).

**Figure 4.**
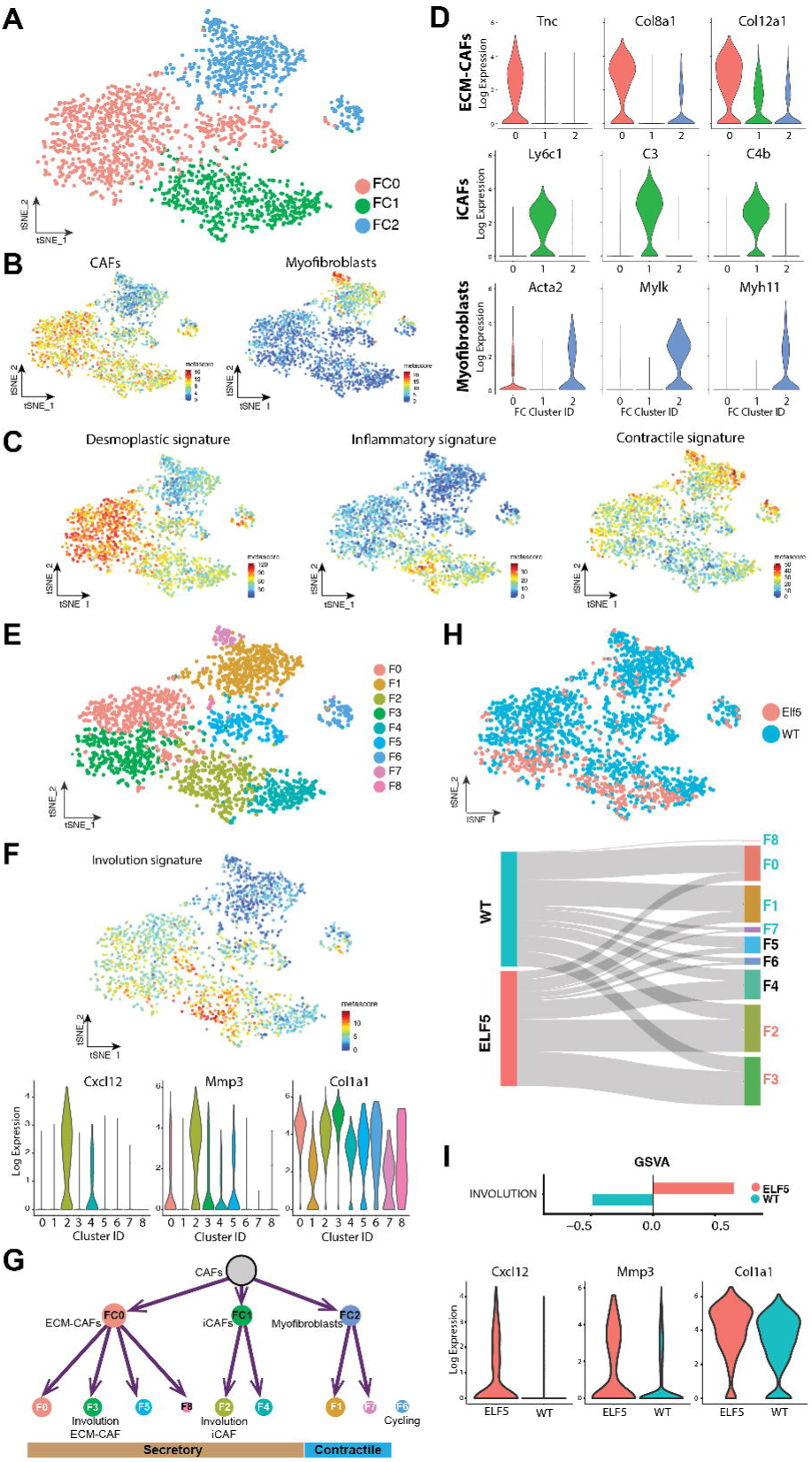
Annotation of the cancer-associated fibroblast diversity in PyMT tumours. **A)** tSNE plot groups defined by k-means clustering analysis showing a total of three cell clusters defined within the fibroblast subpopulation as defined in Figure 1E (prefix FC). **B)** Meta-signatures of the secretory Cancer-associated fibroblasts (CAFs) (upper plot) and myfibroblasts (bottom panel) plotted in the fibroblast tSNE. **C)** Desmoplastic (upper plot), Inflammatory (middle plot) and Contractile (bottom plot) meta-signatures plotted in the fibroblast tSNEs from public data [38]. **D)** Violin plots displaying marker genes for each of the three fibroblast clusters defined in panel A: ECM-CAFs (FC0), immune-CAFs (iCAFs, FC1) and myofibroblasts (FC2). **E)** tSNE plot defined by unsupervised clustering analysis showing heterogeneity of each of the identified fibroblast species. **F)** tSNE illustration of the involution metasignature from [33], bottom panel: violin plots on the nine fibroblast clusters for individual genes from the involution signature. **G)** Cell tree classification of the main lineages of CAFs in PyMT/WT and ELF5 tumours. **H)** Distribution of fibroblasts by genotype (Elf5: red), WT: blue) in the defined tSNE dimensions. Bottom plot: Sankey diagram showing the contribution percentage of each of the genotypes to the cell clusters. Cluster numbers are coloured by the dominant genotype (>2-fold cell content of one genotype), Elf5 (red), WT (green). **I)** GSVA enrichment analysis of involuting mammary fibroblast meta-signatures associated to PyMT/WT and /Elf5 tumours. Bottom panel: violin plots of Cxcl12, Mmp3 and Col1a1 genes in all fibroblasts of each genotype. PyMT/WT (green) PyMT/Elf5 (red).

CAFs are known to be highly diverse, which it was confirmed in our system as classical fibroblast markers [34] failed to be specific to any particular transcriptomically-defined clusters (SuppFig. 6D). Previous unbiased scRNAseq analyses in human tumours has classified two major fibroblast subtypes [35, 36]: activated myofibroblasts, involved in tissue remodelling architecture by physical forces; and secretory CAFs related to ECM synthesis and cyto- and chemokine production [37]. Fibroblast clusters 0 (CF0) and 1 (CF1) were enriched for a secretory CAF signature while the activated myofibroblast signature was concentrated in fibroblast cluster 2 (CF2) (Fig. 4B). Functional annotation using GSVA analysis (SuppFig. 7A) revealed that secretory CF0 were governed by hallmarks of EMT and were enriched for a desmoplastic signature [38], characterised by genes involved in extra cellular matrix (ECM) interactions, suggesting a tissue remodelling function (Fig. 4C and D). Secretory CF1 clearly showed immune-related functions (top enriched hallmarks “Interferon alpha response”, “IL6/JAK/STAT3 signalling” and “Interferon gamma response”) [39], with genes involved in cytokine-cytokine receptor interactions, monocyte recruitment and the complement cascade[38]. CF2 were enriched for a contractile signature, characterised by the expression of genes involved in actin cytoskeleton organisation (Fig. 4C and D).

Taken together, we have functionally characterised three different classes of fibroblasts in PyMT tumours: 2 secretory-CAFs classes, CF0 defined as ECM-CAFs and CF1 defined as inflammatory-CAFs (iCAFs); and a contractile myofibroblast class (CF2). This classification is consistent with the recently reported nomenclature used by Bartoschek and colleagues for CAFs from the PyMT model [40] (SuppFig. 6E and F).

### CAFs from PyMT/ELF5 mammary tumours show features of mimicry involution

Unsupervised clustering of the fibroblasts found in PyMT tumours further identified several subclasses of secretory and contractile fibroblasts (Fig. 4E). Mammary involuting fibroblasts have unique properties that include a high production of fibrillar collagen, ECM remodelling and immune suppression through monocyte recruitment [33]. This specialised class of fibroblast involved in involution are characterised by the expression of Col1a1, Cxcl12, Tgfb1 and Mmp3. We identified a group of ECM-CAFs and iCAFs with high expression of three of these involution markers in PyMT tumours (Col1a1, Cxcl12 and Mmp3, Fig. 4F), clusters F2 and F3). We thus referred these two clusters as involution-iCAFs and involution-ECM-CAFs, respectively. The complete classification of PyMT fibroblasts is shown in Fig. 4G. Involution-CAFs were enriched in PyMT/ELF5 tumours where >2 fold of involution-CAFs, Cluster F2 and F3, belonged to PyMT/ELF5 mice (Fig. 4H). Supplementary Figure 7B shows that such genotype enrichment was not due to batch effects. Consistently, GSVA of the CAF-involution signature and involution marker genes were significantly enriched in the CAFs from ELF5 tumours compared to the WT (Fig. 4I).

The functional annotation of the involution fibroblasts revealed specific differences within the ECM-CAFs or iCAF subpopulations (SuppFig. 7C). For example, the top pathways of the involution ECM-CAFs that were unique of this ECM subgroup, fatty acid metabolism and peroxisome hallmarks, suggested that these cells may correspond to adipocyte-like fibroblasts arisen through a de-differentiation process of mature adipocytes [41] or transdifferentiation process of activated fibroblasts [42]. Adipocyte-derived fibroblasts secrete fibronectin and collagen I and have invasive properties [41, 43]. Moreover, during the stromal remodelling stage in involution, there is a re-differentiation of the adipocytes [32, 44]. Involution iCAFs, on the other hand, showed pathways linked to a wound-healing process, including inflammation through the complement cascade and coagulation [31, 45, 46]. The late stages of involution share many wound-healing properties such as high immune cell infiltration and immune suppression [47], thus these involution iCAFs might be responsible for these two immune-related processes.

Our finding on CAF activation associated with Elf5-alveolar epithelial cells has never been reported before, thus we performed orthogonal validation for the role of CAFs in ECM remodelling by the production of collagen fibres in both PyMT/WT and PyMT/Elf5 tumours. Second Harmonic Generation (SHG) imaging and picrosirius red staining of tumour sections showed an increase in fibrillar collagen coverage in PyMT/ELF5 tumours compared with their PyMT/WT counterparts (Fig. 5A and B). Polarised light microscopy showed that PyMT/ELF5 tumours had a significantly higher proportion of thicker and mature collagen fibres than PyMT/WT tumours (Fig. 5C). Further analysis of matrix ultrastructure revealed a more complex spatial arrangement (peak alignment (+/-10 degrees from peak)) of collagen fibres in PyMT/ELF5 tumours than in PyMT/WT tumours (SuppFig. 8A). At the mRNA level, the proposed markers for involution fibroblasts (Col1A1, Mmp3 and Cxcl12) were highly specific for a subset of cells from the fibroblast compartment (SuppFig. 8B), thus we performed a histological analysis to spatially map and validate these markers at the protein level using immunohistochemistry staining in PyMT/WT and PyMT/Elf5 tumours (Fig. 5D). To further confirm the cellular origin of the collagen deposition, we used a collagen I antibody only recognising alpha-1 type 1 collagen in its monomer form prior fibre assembly, thus labelling the collagen I producing cells. Both COL1A1 and CXCL12 expressing fibroblasts were infiltrated throughout the cancer epithelial foci and organised in clusters, especially in case of PyMT/ELF5 tumours were these cells were also more abundant (Fig 5D and SuppFig. 8C). MMP3 IHC staining shows positive expression in both intra-foci stromal cells and adjacent epithelium but negative expression in epithelial areas away from the stroma, this gradient is consistent with a paracrine effect of MMP3 secretion from invCAFs (Fig. 5D and SuppFig. 8C). The extent of the paracrine gradient was more noticeable in the PyMT/ELF5 tumours compared to the PyMT/WT counterparts.

**Figure 5.**
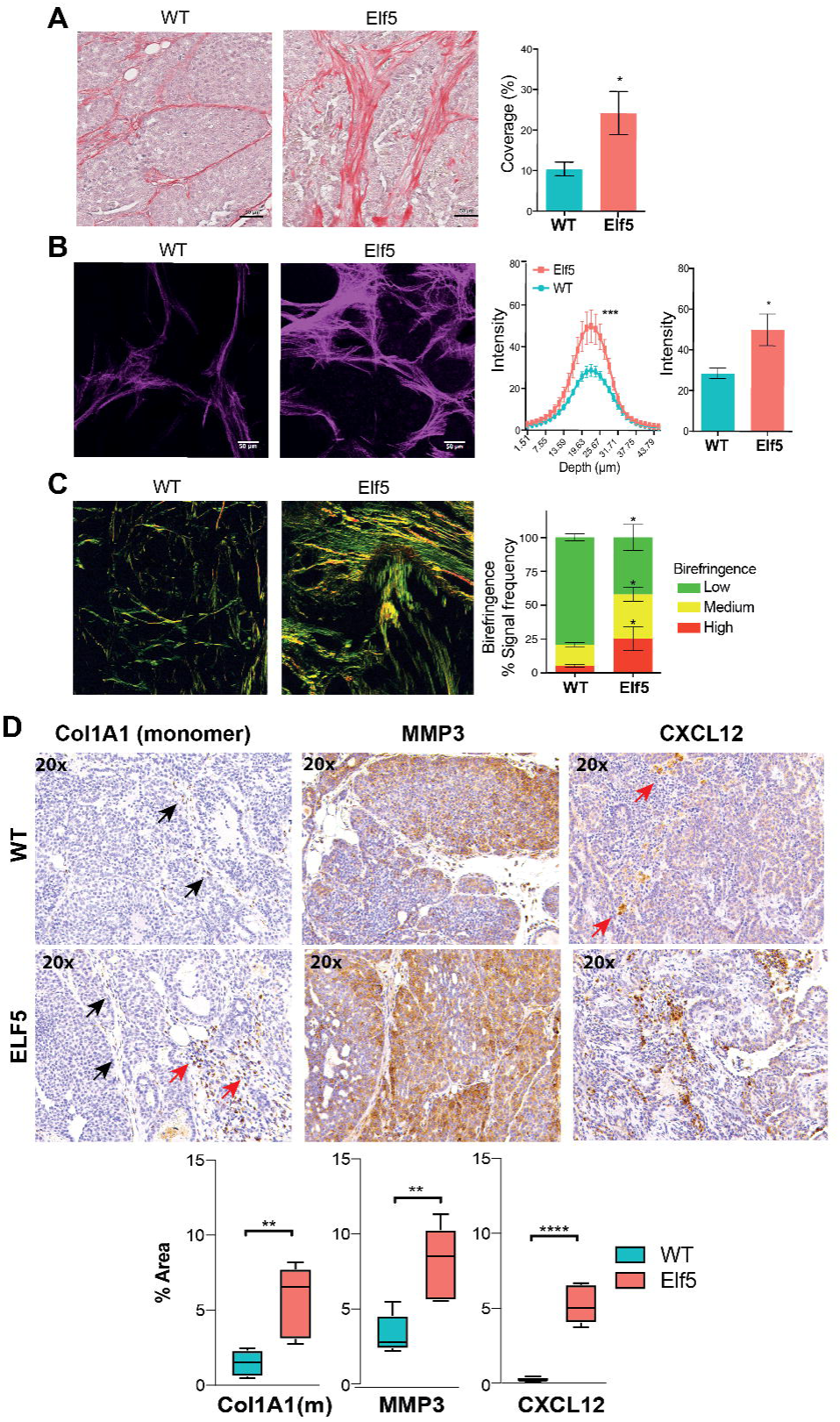
Functional validation involution CAFs in PyMT/WT and /Elf5 tumours. **A)** Representative bright field images and quantification of total coverage of picrosirius red-stained PyMT/WT and PyMT/ELF5 tumours sections n=4 mice per genotype with 10 regions of interest (ROI) per tumour. **B)** Representative maximum intensity projections of SHG signal and quantification of SHG signal intensity at depth (µm) and at peak in PyMT/WT and PyMT/ELF5 tumour sections, n=6 mice per genotype with 6 ROI per tumour. **C)** Polarised light imaging of picrosirius red stained PyMT/WT and PyMT/ELF5 tumour sections, and quantification of total signal intensity acquired via polarised light. Thick remodelled fibres/high birefringence (red-orange), medium birefringence (yellow) and less remodelled fibres/low birefringence (green) n=4 mice per genotype with 10 ROI. **D)** Representative images (top panels) of the immunohistochemistry analysis of COL1A1, MMP3 and CXCL12 in PyMT/WT and /ELF5 tumours and their quantification as % of area stained (bottom box plots). Black arrows show positive staining on elongated cells while red arrows show positive staining in rounded cells. Data corresponds to a n = 5 tumours with at least 5 images/tumour; ** denotes p<0.01, **** denotes p<0.0001

Finally, to confirm the involution-related nature of these markers we performed a parallel histological analysis in a developmental series of the mammary gland, including virgin (V), late pregnancy (18 days *post-coitum*, 18DPC), established lactation (4 days *post-partum*, 4DPP), early involution (1 day involution, Inv. D1) and late involution (4 days involution, Inv. D4) (SuppFig. 9). We found the staining of these three markers in the stroma and in some stages in the stroma in close proximity to the lumen of the mammary gland. MMP3 and CXCL2 followed a very similar pattern of expression, dramatically increasing with pregnancy, dropping during the apoptotic phase of involution and increasing again at late stages of involution. This expression dynamics coincides with time-points of intense active tissue remodelling. The monomeric form of COL1A1 however dropped its expression during pregnancy and remained low until reaching its maximum of expression at late involution. Thus the co-expression of these three involution markers was at its maximum only during the late stages of involution, supporting their association with involution cues and the process of involution mimicry identified in PyMT/ELF5 tumours.

Taken together, these results functionally and histologically characterise invCAFs confirming the correlation between mRNA and protein of their markers, it also corroborates that COL1A1, MMP3 and CXCL12 are useful markers to identify an involution process, and that both the presence and the extent of activation of invCAFs was higher in PyMT/ELF5 tumours.

### Involution signatures predicts for poor prognosis in luminal breast cancer patients in the context of Elf5 expression

High *ELF5* expression is associated with basal breast cancers[18]. We have previously shown that in luminal tumours with higher expression of ELF5 show basal-like characteristics and poor prognosis[19], thus these can be considered as the early steps towards the differentiation into a basal tumour, underscoring ELF5 as a driver of the basal-like phenotype.

We used the METABRIC dataset[56] to investigate whether the involution signature correlated with patient outcome and ELF5 levels. In line with previous reports[18], we found a significant association between high *ELF5* expression and poorer patient prognosis, (SuppFig. 10A). As expected, higher proportion of basal molecular subtype fell into the ELF5-high group, but also a proportion of luminal cancers. Involution was also a predictor of poor prognosis independent of other pathological characteristics such as Ki67, basal subtype and age (<40) (SuppFig. 10A & B). Consistent with our hypothesis, patients with higher expression of *ELF5*, showed significantly higher levels of expression of the involution signature, and this was also confirmed in the TCGA dataset[57-59]. In our PyMT mouse model of luminal breast cancer, our results demonstrate that the acquisition of basal properties driven by ELF5 was associated with the development of tumour progression traits, and in particular the onset of involution mimicry. As shown in Supplementary Figure 8D, in luminal patients, involution predicted for poor prognosis only in the context of high-ELF5 expression. Taking together, these results show that ELF5 expression is a useful marker to identify luminal patients that present poor outcome associated to an involution process.

### Characterisation of the cell-to-cell interconnections involved in the cancer-associated involution mimicry

Involution is a multicellular process, thus we used CellphoneDB to identify networks of cell-to-cell communication within PyMT tumours [48]. CellphoneDB analysis in PyMT tumours revealed an intricate network of cell interactions. Figure 6A shows the interactome of all cell clusters defined in this study (Figs. 1, 2 and 4, summarised in SuppFig. 7D). These interactions were ranked from strongest to weakest (count above mean 0.3) which allowed us to categorise them in three levels: 1) fibroblast-fibroblast interactions were the strongest, followed by fibroblast-epithelium connections; 2) endothelium-fibroblast or -epithelium and epithelium-epithelium interactions were ranked as mid interactions; and 3) the weakest connections were found in the immune cell compartment. A complete classification of the predicted cell-to-cell interactions can be explored using our html-based interactive tool (https://odonoghuelab.org:8000), where cell types are grouped by unsupervised clustering of common signalling pathways. These results show a pivotal role of CAFs as hubs of communication within the TME.

**Figure 6.**
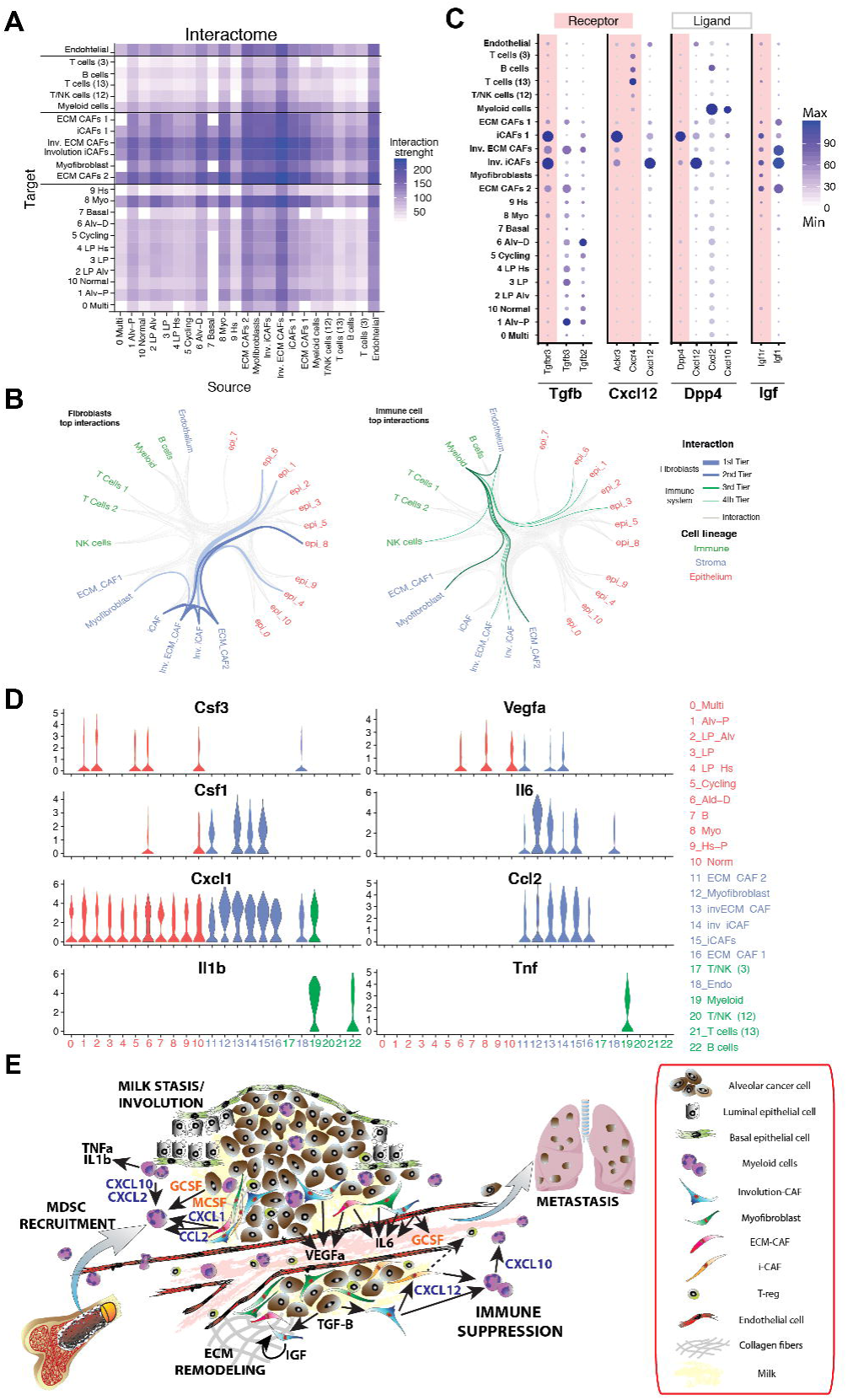
Interactome of PyMT tumours. **A)** Heatmap of the cell-cell interactions of all cell types from PyMT tumours based on CellphoneDB. Cell classification is based on the global annotation shown in SuppFig. 7D. For simplicity, F6 and F8 were removed and F1 and F7 (myofibroblasts) were pooled together as a single cluster. The scale at the right-hand side shows the interaction strength based on the statistical framework included in CellphoneDB (count of statistically significant (p<0.01) interactions above mean= 0.3, see extended methods). **B)** Graphical representation of all significant cell-cell interactions identified by CellphoneDB using the parameters of more than 10 significant interactions with a mean score greater than 0.3, number cut as more than 10 connections and number split 10 (all lines in all colours included grey lines). Red colour corresponds to the cell types from the epithelial compartment; blue colour represents the cells from the stromal compartment and the green colour are the cells from the immune compartment. Different number splits were applied to establish the most significant interactions for the fibroblast and immune cell types. Fibroblast showed the strongest interactions (highlighted as blue lines) when a number split of 67 (1^st^ Tier) and 50 (2^nd^ Tier) were used. The immune system showed weaker interactions (highlighted as green lines) at a number split of 15 (3^rd^ Tier) and 11 (4^th^ Tier). For an interactive version of this representation visit https://odonoghuelab.org:8000 **C)** Representative dot plots of ligand (no background)-receptor (red background) pairs. The size of the circles is relative to the number of cells within each annotated cluster that showed a positive expression of each gene and the blue gradient represents the average scaled expression. **D)** Violin plots of genes from canonical pathways known to recruit and expand MDSCs in all cell cluster as defined in SuppFig.7D. Colour code of the cell clusters: Red: epithelial compartment. Blue: stroma and Green: Immune cells. **E)** Proposed molecular model of involution mimicry driven by Elf5 expression in the epithelial compartment where CAFs and MDSCs are the major cell types involved.

Involution accounts for an interactive cell network where epithelial cells, fibroblasts and immune cells are the key players [32]. We found that when the most restrictive cut offs were applied (Fig. 6B, 1^st^ and 2^nd^ Tier, blue lines), the ECM-CAFs (involution and ECM CAFs 2, which correspond to FC0 (defined in Figure 4) and involution iCAFs showed the largest number of interactions between themselves and also with the two Elf5 enriched-alveolar epithelial cells clusters (epithelial Clusters C1 and C6 defined in Figure 2), the myoepithelial cells and the luminal progenitor hormone sensing cells (LP Hs, epithelial cluster C4). In the immune system interactive network (Fig. 6B, 3^rd^ and 4^th^ Tiers, green lines), the myeloid compartment showed the highest number of connections and interestingly, myeloid cells communicate with the two alveolar epithelial clusters, with ECM CAFs (involution and ECM CAFs 2), involution iCAFs, myofibroblasts and with the endothelium. Involution is associated with the M2-like innate response, thus these results are in line with an activated and interactive innate myeloid response [9, 31, 49].

We then investigated what particular ligand/receptor connections were found among CAFs, alveolar epithelial and myeloid cells in order to find common pathways linked to involution (Fig. 6C). Overall, we found pathways associated with the multi-functional aspects of involution (TGFβ axis) [50, 51]; networks associated with an immune suppressive ecosystem (Cxcl12 and Dpp4 axis) [33, 52] and ECM remodelling (IGF axis) [53]. Transforming growth factor β2 and 3 (TGFβ2 and TGFβ3) are ligands produced by CAFs and the two alveolar epithelial clusters, while their receptor TGFβR3 is exclusively expressed in involution ECM and iCAFs and in iCAFs-1 (corresponding to FC4). The chemokine Cxcl12 is exclusively expressed in the involution iCAFs and two of its known receptors, CXC chemokine receptor 4 (CXCR4) and atypical chemokine receptor 3 (ACKR3 or CXCR7), are expressed in iCAFs-1 and B and T cells, respectively. Dipeptidyl peptidase-4 (Dpp4) expression was restricted to iCAFs-1 and showed a highly interactive axis; Dpp4 connected with the involution iCAFs through Cxcl12 and with myeloid cells through Cxcl2 and Cxcl10. The IGF1/IGF1R signalling pathway was found to be highly specific to the two involution CAF subtypes, ECM and immune, in which a predicted positive feedback loop among these cells was observed. Of note is that IGF signalling is classically associated with collagen and ECM remodelling in fibroblasts [53].

The myeloid cell compartment is also intimately associated with the process of involution. We have previously demonstrated that tumour infiltrated myeloid-derived suppressor cells (MDSCs) are involved in the formation of lung metastasis in PyMT/ELF5 alveolar tumours[19]. As tumour infiltrated myeloid cells constitute a low abundance population[54] we kept it as a unique cell cluster, however, our scRNAseq analysis was able to map the cellular mechanism for MDSC expansion, recruitment and malignant activation previously described in these alveolar tumours[55], identifying the cell of origin of well-known signalling pathways involved in this process (Fig. 6D). G-CSF (Csf3), a potent growth factor that attracts and recruits MDSCs, was expressed in cancer cells associated with the alveolar clusters, which would explain the increased infiltration of MDSCs in PyMT/ELF5 tumours[19]. Malignant activation of MDSCs is also driven by M-CSF (Cfs1), mainly expressed by CAFs. In addition, VEGF (Vegfa), was highly secreted by alveolar, basal cancer cells and CAFs, and IL6 is mainly secreted by CAFs and endothelial cells, these are factors that contribute to MDSC expansion. Potent MDSCs chemo-attractants were expressed by a diverse range of cell types within the tumour; Cxcl1 is ubiquitously expressed in most cell types, while Cxcl2 was highly expressed in the myeloid cell cluster, suggesting this is a mechanism that tumours may use to maintain a feedback loop enabling continuous myeloid cell recruitment and sustained inflammation. Other recruitment factors such as Cxcl12 and Ccl2 are specifically secreted by CAFs. Altogether, our results show that MDSCs recruitment and malignant activation is not just driven by cancer cells but by a complex molecular network orchestrated by alveolar cancer cells and involving multiple cell compartments in the TME, mainly CAFs, endothelial cells and tumour infiltrated myeloid species.

In summary, we have found that cells in the TME of PyMT tumours are heavily involved in the promotion of tumour progression. We have characterised the intercellular molecular pathways originated in alveolar cancer cells that resemble a deregulated established involution. This involution mimicry results in an inflammatory process characterised by malignant CAF education and recruitment of MDSCs, which ultimately results in the acquisition of functional traits of tumour progression (Fig. 6E).

## Discussion

The MMTV-PyMT model gives rise to mammary tumours of luminal progenitor origin as classified by conventional transcriptomics[13, 60]. Our findings however underscore the expansive plasticity of the luminal progenitor compartment identifying “malignant states” associated with the mammary epithelium cell lineage specification. Our results unite tumour heterogeneity with developmental cues and highlight the intrinsic capacity of tumours to develop distinct molecular subtypes, supporting the current understanding of progenitor cells as the cell of origin of basal and luminal cancers (reviewed in [12])

Here we study the functional consequences of the specification of the ER-basal subtype of breast cancer through the differentiation from the luminal progenitor cells towards the pregnancy-derived alveolar lineage[4, 61]. ELF5-specified alveolar cancer cells subsequently activate cell-extrinsic mechanisms within the TME reminiscent of mammary morphogenesis, which ultimately was associated with the acquisition of tumour progression cues, including increased immune infiltration of immunosuppressive MDSCs, vascular leakiness and angiogenesis, hypoxia and collagen deposition, which are all hallmarks of more aggressive tumours and are linked to metastases, as we observe in our Elf5/PyMT model and similar to that induced by pregnancy[19]. Interestingly although hypoxic conditions are commonly associated to undifferentiated tumours[62], alveolar tumours raised from a differentiating mechanism induced by Elf5 are extensively hypoxic, this particular feature might be also a reminiscence of lactation in the normal mammary gland, where hypoxic conditions emerge to accommodate the intense metabolic activity of milk production[63].

Clinically, patients with luminal tumours receive standard-of-care anti-estrogen therapy, thus adoption of the alveolar fate provides a route of escape from endocrine therapy, as these cells are insensitive to estrogen[18] and subsequently give rise to a basal-like tumour[18] with much poorer survival rate[19]. We have found that expression of *ELF5* is characteristic of luminal cancers that are in transit to a basal-like subtype, thus breast cancer patients that show higher Elf5 expression activate involution mimicry pathways that are associated with poor prognosis. In addition, the pro-metastatic effects of Elf5 are concomitant with mimicry of mammary involution. In this line of evidence, involution has been previously shown to be involved in post-partum associated breast cancer (PABC)[17] however the molecular drivers of this process had not been not fully identified.

Our Elf5-restricted MMTV-PyMT model reproduces the particular effect of pregnancy on the specification of the alveolar cell lineage in the context of cancer, shedding light upon the molecular mechanisms involved in these effects of pregnancy in breast cancer. Mammary involution is a wound-healing-like process which includes tissue remodelling with fibroblast activation to reorganise the ECM and accommodate the epithelial reduction, and myeloid infiltration to remove all cell fragments[8, 9]. Our data presented here supports the hypothesis that the induced expression of *Elf5* stimulates the production of milk by alveolar cells, which in turn accumulates and triggers an inflammation-mediated TME remodelling associated to aberrant involution. This mimicry involution process results in alveolar cell death that is rapidly restored by ELF5-driven differentiation of luminal progenitors and a chronic induction of inflammation by involution-iCAFs, characterised by the activation of STAT3/JAK and IFNgamma pathways[39] and myeloid cells encompassing ECM remodelling driven by involuting ECM-CAFs (Fig. 6E). We have focussed this study in the luminal MMTV-PyMT mouse model of cancer; thus, further work is necessary to assess the potential benefit of targeting involution mimicry as a strategy for generalised anticancer therapy.

Our scRNAseq analyses have also allowed us to identify putative molecular candidates responsible for this epithelium-fibroblast-myeloid crosstalk during mimicry of involution (Fig. 6). We find that only alveolar epithelial cells express TGFβ2 and 3 ligands and their receptor TFGβR3 is expressed in involution CAFs. TGFβ is linked to tumour malignancy by affecting both cancer cells and the TME, including CAF activation and immune suppression[50] suggesting that this could be a specific pathway driving mimicry involution. In fact, TGFβ3 is induced in response to milk stasis during the first stage of involution for the induction of alveolar apoptosis and TGFβ signalling is also key in the second stages of normal involution for the induction of ECM deposition and immune suppression[51]. Altogether, this suggests that TGFβ signalling might be one of the pivotal pathways triggered by Elf5-alveolar cells and thus a potent pathway to be therapeutically targeted.

We also find specific pathways of crosstalk among involution CAFs, other CAF-types and the tumour-infiltrating immune system, which might be activated following this ELF5-mediated TGFβ activation. These include the expression of Cxcl12 by the involution-iCAFs and IGF1 from both ECM and inflammatory involution CAFs. Cxcl12 is a stromal chemokine that attracts leukocytes through CXCR4[64] and enhances cell adhesion and survival through ACKR3[65], these receptors are expressed in T/B cells and iCAFs, respectively. Interestingly, involuting mammary fibroblasts that highly express Cxcl12 induce monocyte recruitment and are associated with blockade of CD8+ T cell tumor infiltration[66]. Cxcl12 is also a known recruitment factor for MDSCs[55]. In this context, CAFs also show high expression of other classical markers for MDSC recruitment and activation, including M-CSF, IL-6 and Ccl2[55] underpinning the high infiltration and activation of MDSCs previously described in Elf5-high tumours[19]. Dpp4 is a ligand from one of the iCAFs groups that could also build the communication bridge between Cxcl12-expressing involution iCAFs and Cxcl2 and Cxcl10-expressing myeloid cells. Dpp4 cleaves and inactivates these chemokines promoting an immunosuppressive environment by the inhibition of the recruitment of effector T cells[67] and by the promotion of chronic inflammation[68]. This also suggests that the M1-innate response from myeloid cells and fibroblasts is inhibited by Dpp4-expressing iCAFs. In addition, Costa and colleagues recently demonstrated that a subtype of CAFs from human breast tumours was responsible for the increase of regulatory T cells (Tregs) through the expression of Dpp4[52], so the involvement of Tregs in Elf5-mediated immunosuppression might be another area to further explore. Thus, our data suggests that most of the immunosuppressive signals come from the presence of activated MDSCs in the tumour, mainly recruited by involution CAFs signaling as a response to the involution mimicry process initiated by alveolar cancer cells (Fig. 7E).

Altogether our data illustrates a highly interactive TME, particularly with CAFs and myeloid cells, with cancer-differentiated cells that are pivotal initiating the events that result in tumour immunetolerance and metastasis concomitant with mimicry involution. Our data highlights the relevance of targeting cancer-associated cell species as a strategy for anticancer therapy an approach particularly important in the context of PABC.

## Conclusions

- Drop-seq resolves tumour cell diversity of PyMT mammary tumours revealing strong cancer cell heterogeneity.
- Cancer epithelial cells are hierarchically organised and resemble the cell lineages in the mammary gland. We show that each cell lineage has associated specific hallmarks of cancer progression; despite PyMT mouse model being transcriptional classified of luminal-origin, we reveal the presence of all mammary epithelium lineages and their intermediate progenitors within these tumours.
- Elf5 drives alveolar lineage differentiation in PyMT tumours and redefines the tumour cell diversity, resulting in the acquisition of novel cancer cell states towards basal-like characteristics concomitant with the acquisition of metastasis.
- We have identified a multicellular mechanism of tissue morphogenesis associated to unresolved lactation, involution mimicry involution mimicry, as an aberrant developmental mechanism associated to Elf5 high tumours, both in mice and humans, during the luminal to alveolar switch.
- A subpopulation of secretory CAFs (involution ECM and involution iCAFs) from the TME play a pivotal role during Elf5-associated involution mimicry, characterised by a highly interconnected TME with cancer cells.
- ELF5 could be used as a biomarker to identify luminal patients that present poor outcome associated to an involution process.

## Materials and Methods

### Animal statement, doxycycline induction

All mice of this study are in a pure FBV/n background with more than 20 generations of backcrossing. To promote tumour progression, ELF5 expression was specifically induced in mammary epithelial cells since puberty (6 weeks-old) through doxycycline containing food as performed previously[19]. At ethical endpoint (7-13% tumour/body weight) mice were euthanized and tumours harvested and processed for single cell digestions as previously performed[19]. All experiments involving mice have been approved by the St. Vincent’s Campus Animal Research Committee AEC #14/27, #17/03 and #19/02.

### Tumour digestion and FACS

Mammary tumours were collected from MMTV-PyMT mice at 14 weeks of age, when tumour weights were 10±3% of total weight and digested to single cell suspension as performed before^15^ (also see Extended Methods). Flow cytometry analysis were performed on BD FACSymphony– High-Speed Cell Analyser or the FACSAriaIII (Becton Dickinson) following automated compensation (FACSDiVa) using a 100um nozzle. Unspecific binding was prevented using the ells were stained using the BD Pharmingen Purified Rat Anti-Mouse CD16/CD32 (Mouse BD Fc Block™) (Clone 2.4G2) and Rat IgG (Jackson ImmunoResearch Rat Gamma Globulin in a buffer containing DNase I (Sigma-Aldrich). Subsequent immunostaining was performed using the following flourescently tagged antibodies: CD45R/B220 (clone RA3-6B2), CD31 (clone MEC 13.3), CD8a (clone 53-6.7), Ly-6G (clone 1A8) and CD4 (clone GK1.5) from BD Biosciences; F4/80 (clone BM8) and CD11b (clone M1/70) from eBioscience; and CD45 (clone 30-F11), CD3 (clone 17A2), CD326 (Ep-CAM) (clone G8.8) and Ly-6C (clone HK1.4) from BioLegend. Data was analysed using FlowJo software (version 10.4.2).

### Auto macs cleanup

The viability of the cells was assessed by FACS. All tumours that contained less than 80% alive cells were labelled with Annexin specific MACS beads and dead cells were removed by passing the labelled cells through the autoMACS® Pro (Miltenyi) (see more details at [69]). A preparation of cells containing a high proportion of viable cells (>85% viability assessed by DAPI in FACS) were loaded into the microfluidic Drop-seq pipeline.

### Drop-seq

Cells were captured using the microfluidic devices described for Dropseq [24] following the Online Drop-seq Laboratory Protocol version 3.1 (www.mccarrolllab.com/dropseq). The tagmented and multiplexed cDNA libraries were sequenced in Nextseq 500 using Nextseq 500 High Output v.2 kit (75 cycles, Illumina Cat# FC-404-2005) following Macosko et al. recommendations with the following modifications. Drop-seq denatured libraries were loaded at 1.3pM final concentration and were spiked in with 10% of 1.8pM PhiX Control v3 (Illumina Cat# FC-110-3001). Sequencing specifications were as follows: 26bp Read1, 51bp Read 2 and single Nextera indexing (8bp). A total of ∼3,000 cells/ per run were sequenced.

### Bioinformatic analysis (see Extended Methods and SuppFig. 1)

#### Multiphoton microscopy

Second Harmonic Generation (SHG) signal was acquired using a 25x 0.95 NA water objective on an inverted Leica DMI 6000 SP8 confocal microscope. Excitation source was a Ti:Sapphire femtosecond laser cavity (Coherent Chameleon Ultra II), tuned to a wavelength of 880 nm and operating at 80 MHz. Signal intensity was recorded with RLD HyD detectors (420/40 nm)). For tumour samples, 5 representative regions of interest (512 μm x 512 μm) per tumour were imaged, for CDMs 3 representative areas of 3 technical replicates over a 3D z-stack (20 μm depth; 20 μm depth for CDM, with a z-step size of 1.26 μm). SHG signal coverage in tumour samples was measured with ImageJ (National Institutes of Health, Bethesda, MD, USA). For CDMs mean SHG intensity was measured using Matlab (Mathworks). Representative images of maximum projections are shown.

Blood vessel patency analysis was performed injecting 10 ul of quantum dots blood tracers (655nm Life Technologies) through the tail vein of the animals **(Supp Videos 1 and 2)**. Images were acquired with a 25x NA0.95 water objective. A dichroic filter (560 nm) was used to separate the GFP signal from quantum dot emission, which were further selected with band pass filters (525/50 and 617/73, respectively).

#### Western Blot and Immunohistochemistry

Western Blot analyses of protein were performed as previously detailed in [18]. The primary antibodies not described in Kalyuga et al were used as follows: E-Cadherin (BD bioscience #610181, 1:10,000) and Vimentin (Leica Biosystems # VIM-572-L-CE). Mouse mammary glands from FVB/N mice along different stages of the mammary gland development and tumours from PyMT/WT and PyMT/ELF5 mice were fixed, defatted, dehydrated, stained in carmine alum and caraffin embedded as previously described[70]. Sections were stained with the rabbit anti-mouse milk proteins antibody (Accurate Chemical & Scientific CO, # YNRMTM) diluted 1:12,000 as per[70]. Staining with COL1A1 (E6A8E), CD8 and CD4 (Cell Signalling Technlogies); and MMP3, CXCL12/SDF1 and SMA (Abcam) were performed using the Leica Bond RX. Staining with CD45 (BD Pharmigen) was performed using the DAKO autostainer instrument. Image quantification was performed as previously described[71]

#### Polarized light microscopy of picrosirius red stained sections and quantification of collagen content and birefringence

Paraffin-embedded samples were cut into 4 μm sections and stained with 0.1% picrosirius red for fibrillar collagen according to manufacturer’s instructions. As previously performed [72, 73] polarized light imaging was performed on a Leica DM6000 fitted with a polarizer in combination with a transmitted light analyser. Quantitative intensity measurements of fibrillar collagen content and birefringent signal were carried out using in house scripts in ImageJ. The relative area of red-orange (high birefringent) fibres, yellow (medium birefringent) fibres, and green (low birefringent) fibres (as a % of total fibres) was calculated.

#### Hypoxia analysis

Tumours and lungs were harvested and fixed in 10% neutral-buffered formalin overnight. FFPE tissues were sectioned to 4µm for IHC staining as described in[19]. Following the manufacturer’s instructions, the FITC-conjugated pimonidazole primary antibody (Hypoxyprobe, 1:1000) and the anti-FITC secondary antibody (Hypoxyprobe 1:100) was used to stain for the pimonidazole adducts. Quantification of hypoxia in IHC was performed using Andy’s DAB Algorithm [71].

## Declarations

### Ethics approval

All experiments of this study involving mice have been approved by the St. Vincent’s Campus Animal Research Committee AEC #14/27, #17/03 and #19/02.

### Availability of data and materials

The datasets generated during the current study will be available in the GEO repository upon publication. A confidential reviewer link can be facilitated upon request.

### Authors’ contribution

Conceptual and experimental design: FVM and DGO. Data acquisition and experimentation: FVM, RS, AMKL, LC, MP, LRF, ZK, DGO. Data analysis: FVM, AMKL, MP, LRF, KJM, AM, NF, TRC, DGO. Provided reagents and expertise: JRWC, SRO, TRC, CJO. Bioinformatic Analysis: FVM, BG, DLR, YCS and DGO. Data visualisation: FVM, BG, NS, SIO and DGO. Data interpretation: FVM, BG, PT, CJO and DGO. Manuscript writing: FVM and DGO. All authors read and approved the final manuscript.

## Acknowledgments

We thank Dr Marina Pajic and Dr Christine Chaffer from the Garvan Institute for their review and feedback on the manuscript. We also thank the Histopathology Facility from the Garvan Institute for their IHC services.

