## Supplementary Information for "Single cell transcriptomics reveals involution mimicry during the specification of the basal breast cancer subtype"

**Functional heterogeneity of PyMT tumours is associated to lineage origin (cont).**

Cluster 0 (C0) was consistent with undifferentiated cells or uncommitted progenitors (depleted Notch signalling). C0 was characterised by a high enrichment on proliferative and cell cycle associated hallmarks suggesting that these could be a cell of origin of a highly aggressive cancer. Interestingly C0 was enriched on inflammatory associated hallmarks, such as IL6/Stat3, Interferon a and g, inflammatory response, allograft rejection and ROS. This cluster is consistent with immunogenic tumours such as those of the TNBC subtype.

The luminal progenitor group was formed by clusters Cluster 2, 3 and 4. Cluster 2 was functionally similar in both genotypes with a slight cell enrichment in Elf5 tumours. C2 was characterised by a profound depletion on differentiation pathways such as Notch and TGFβ as well as proliferative and metabolic pathways, this suggests that C2 has a highly multipotent progenitor profile with high epithelial score (depletion of EMT), consistent with a transitory amplified progenitor. C3 was characterised by the expression of genes signatures associated to IFN α and γ, this suggests that through the differentiation to the specific lineages and, in particular, in the alveolar compartment, IFNα and g signalling gets progressively lost. For example, C0 presented enrichment for IFNα and g, this is conserved in the luminal progenitor cells C3 and only partially conserved in C1 (IFNα enriched and IFNγ depleted) to end up negatively both enriched in C6 and poorly enriched in C9. Interestingly, C3 as a luminal progenitor was depleted for signatures associated to hormone response (androgen, estrogen response early and late, IL2/Stat5) as well as hallmarks associated with TME communication (e.g. hypoxia, coagulation, angiogenesis) and inflammation (inflammatory response, ROS, complement) and a markedly low EMT, proliferative and differentiation pathways. This cluster was consistent with a non-committed luminal progenitor identity, with no clear commitment to either alveolar or the hormone sensing lineages. Albeit a slight enrichment in WT tumours, the functional activity of these cells is very similar between genotypes indicating a true luminal progenitor identity. Amongst the few differences that could be found between C3_WT and C3_Elf5 cells, it stood out an increase in the score for the signatures related to Notch and TGFβ, this could be consistent with a bias towards a transition to a more differentiated state in Elf5 tumours.

The functional annotation for Cluster 4 is consistent with the previous results of marker genes and trajectory analysis shown in Figs 3 and 4, placing this cluster as a luminal progenitor committed towards the hormone-sensing lineage. Cluster 4 was heavily enriched in WT tumours where Hormone Sensing differentiated cells (C9) are much more frequent. LPs are characteristically depleted in hormone signalling hallmarks, however C4 scores for hormone related hallmarks (ER_early [-0.110]; ER_late [-0.128]; Androgen [-0.017]) were comparatively higher than C2 (ER_early [-0.-0.323]; ER_late [-0.171]; Androgen [-0.164])and C3 (ER_early [-0.209]; ER_late [-0.168]; Androgen [-0.238]), consistent with their transition towards a Hs lineage but still retaining the functional characteristics of a luminal progenitor. Similarly to the other clusters, C4 Elf5 cells presented high Notch signalling enrichment score compared to C4 WT.

Consistently with previous studies, Elf5-driven cell clusters associated to the alveolar lineage, clusters 1 and 6 (C1 and C6), presented a negative enrichment on estrogen responsive genes, which is presumably associated to the opposing transcriptional force that Elf5 drives on ER signalling [1, 2]. Similarly, a downregulation of genes involved in EMT were characteristics of these two clusters [1, 3]. These clusters were also characterised by an enrichment of the IL6/Stat3 signalling pathway, a pathway associated with tumour progression and metastasis but also with induction of cancer immunetolerance [4], and cholesterol homeostasis, also associated with breast cancer progression, inflammation and metastasis. Interestingly aspects of innate immune related pathways (Hallmark Complement) were highly enriched in both alveolar clusters. Reduced cell cycle progression and cell proliferation has been associated to Elf5 expression [2] and tumour growth [1] this is consistent with the observed decline of Hallmarks related to cell cycle progression (G2M, E2F, mitotic spindle and Myc targets) and cell growth and proliferation (i.e Hedgehog). All these characteristics have been previously associated to Elf5 rich tumours including in the PyMT mouse model [1]. In addition, C6 was characterised by enriched genes in response to hypoxia, coagulation and involved in angiogenesis, suggesting a strong sensitivity of these cells to signals of the extracellular ecosystem, as Elf5 tumours have been shown to exhibit strong angiogenesis, and vascular leakiness [1]. Similarly, cells from C6 seemed to be responding to oxidative stress (ROS and Xenobiotic signalling pathways), this is associated with the formation of chronic inflammation and recruitment of leukocytes, and the depletion on IFN signalling in this cluster might be associated to increase of immune tolerance and recruitment of immunosuppressive leukocytes, something that has been previously shown in this model associated with Elf5-high tumours [1].

The hormone sensing linage was represented by Cluster 9. This cluster was characterised by an enrichment of a number of hallmarks associated to hormonal response, Estrogen, (late and early), androgen and IL2/Stat5 pathways. Interestingly and similarly to what it was observed in C6, these more differentiated cells in both lineages were strongly associated to communication with the TME, including hypoxia, angiogenesis and inflammation associated hallmarks.

The basal/myoepithelial subtype was formed by clusters 7 and 8. Cluster 7 (C7) is consistent with cells from the myoepithelial lineage. These cells were heavily involved in inflammation-associated hallmarks (complement, allograft rejection, inflammatory response, etc.) and hallmarks associated with TME interactions such as hypoxia and angiogenesis. Such characteristics were also found in the undifferentiated cells from C0, and often undifferentiated multipotent cells and myoepithelial cells showed similar gene signatures. However, in direct contrast with cells from C0, C7 cells present a strong mesenchymal phenotype, characterised by EMT and a higher myogenesis score, and an enrichment in Notch signalling, indicating restriction of stem cell activity and commitment towards a more differentiated fate.

Cluster 8 was consistent with a basal/ myoepithelial lineage and cells of this lineage were more frequent in WT tumours, presumably by the luminal skew produced by the Elf5 differentiation force. Cells that belong to this cluster presented enrichment for Notch and TGFb pathways, indicating a differentiated stage, however these cells are rich in EMT markers, typical of a myoepithelial lineage. Amongst the differences by genotype, a negative enrichment of innate immune inflammatory properties (complement, IFNg, and oxidative stress signals) as well as reduced proliferative hallmarks (mitotic spindle, G2M checkpoints and E2F targets) in Elf5 cells compared with WT cells were the major highlights (Supplementary Fig. 7, Heatmap).

### Extended Methods

#### Tumour digestion

Areas of necrosis were excluded and only regions of epithelial mammary carcinomas were used for digestion and analysis. Tumours were manually dissected into 3-5mm pieces using a surgical scalpel blade before being chopped to 100µm with maximum blade force on a McIlwain Tissue chopper. Tumour samples were incubated with 15,000 U of collagenase (Sigma #C9891) and 5,000 U of hyaluronidase (Sigma #H3506dissolved in DMEM high glucose with 5% FBS at 37°C shaking in 220rpm for 1 hour. Samples were briefly disrupted with a pipette every 15 min during the incubation time to ensure the tissues were sufficiently resuspended. 0.25% trypsin (Gibco #15090-046) in 1mM EGTA was used to digest the samples for 1min in 37°C waterbath before proceeding with a 5 minute incubation with 5mg/mL of dispase (Roche #04942078001) dissolved in PBS in 37°C waterbath. Red blood cells were then lysed with 0.8% ammonium chloride (Sigma #A9434) dissolved in water for 5 min in 37°C waterbath. Samples were washed with PBS containing 2% FBS and spun at 1200rpm for 5min at 4°C between each step. The supernatant was aspirated and 1mg/mL DNase (Roche) was mixed with the sample before incubation with each step. Finally, cells were filtered through a 40µM nylon mesh (Corning) and resuspended in PBS with 2% FBS. FACS was performed as detailed in [1, 2].

#### Bioinformatic analysis

The sequencing output was analysed using the McCaroll lab core computational protocol with a custom genome (mm10 plus Trinity assemblies of transgene sequences [5]) and gene annotation (gencode vM14 plus). Seurat (v Seurat_2.3.4 [6]) was the main platform for downstream analysis.

A total of 26,613 cells were sequenced (18,828 from WT tumours and 7,785 from ELF5 tumours) (SuppFig 2A). Firstly, we removed low-quality cells by modelling mitochondrial to nuclear gene content to <15% [7] and considering differences between homeostatic tissues and tumours. We subsequently removed outlier cells that contained more than 4,000 genes as they could potentially constitute cell doublets (SuppFig. 2B). Thus, DGE matrices were trimmed for quality metrics (>200 genes, <15% mitochondrial genes, and identified genes expressed in at least 3 cells), as a result 15,702 high quality cells (11,490 PyMT/WT and 4,212 PyMT/ELF5) with a total of 28,945 genes proceeded with downstream analysis. A total of 6,176 informative genes were identified based on expression and variance and organised into principal components. Two thirds of the total variation of the system was defined by the first 20 PCs. Downstream analysis was performed according to Butler et al with UMI number regression and 20 principal components of variable genes being used for dimensional reduction (tSNE) and cluster calling [6] (SuppFig 2C). Elf5 expression was restricted to the PyMT/ELF5 tumours (SuppFig.2D). PyMT tumours are derived from the signal of the Polyoma middle-T oncogene in congenic FBVn animals of pure background, thus the effects of the environment are minimised as the individuals are exposed to identical controlled standard laboratory conditions.

Clustering tree [8] was employed in conjunction with cluster validation, and split error estimation across a range of clustering resolutions to identify optimal resolution values. CCA was performed on experiments split by genotype using the union of the top 2000 variable genes for each genotype. No cells were discarded from downstream analysis to retain unique subpopulations between the experiments and downstream analysis was performed as above using CCA in place of PCA.

Monocle (v2 ole [9]) was used to assemble cells assigned to epithelial clusters along a pseudotime vector generated from single cell expression of "Gorsmy" [10]. States were assigned using DDRTree according to the manual.

CellphoneDB [11] (www.CellPhoneDB.org) was performed for 100 cells expressing the highest number of genes in each cluster. A specific interaction was considered as significant if $p<0.01$ and mean score $>0.3$. Expression values of ligand/receptor gene pairs were plotted using Seurat DotPlot function for all cells in each cluster.

GSVA [12] was calculated for averaged expression values for clusters or Poisson distributed counts data using gene mouse hallmark gene lists downloaded from http://bioinf.wehi.edu.au/software/MSigDB[13, 14]. Metasignatures were generated by calculating the sum of scaled expression scores for all genes nominated for the signature in each cell. The relative contribution of each gene to the score was used to rank genes in the signature.

Clinical samples and survival analysis

We used the METABRIC (Molecular Taxonomy of Breast Cancer International Consortium)[15] dataset and the TCGA (The Cancer Genome Atlas) dataset[16-18] for our survival analysis. These datasets contains RNAseq and microarray data as well as detailed clinical information from breast cancer patients. Data was accessed using cBioPortal (https://www.cbioportal.org/) [19, 20].

Patient survival analysis was performed using the "survminer" package (https://github.com/kassambara/survminer). Survival Curves were drawn using 'ggplot2'. R package version 0.4.3 on METABRIC data accessed from the R-Based API for Accessing the MSKCC Cancer Genomics Data Server, CGDSR (R package cgdsr version 1.2.10). Cohorts were split by ELF5 expression and then by metascores and overall survival was compared by cox proportional hazards analysis.

**Supplementary Figure legends.**

**Supplementary Figure 1. A)** Schematic representation of the transgenic models used in this study, MMTV-PyMT and mammary restricted (MMTV) doxycycline-inducible (rtTA) Elf5 expression mouse**. B)** Images of histological sections of lungs from 14 week-old PyMT/WT and /ELF5 mice and the quantification of the area covered by metastases (n=5). **C)** Representative pictures of immunofluorescence of blood vessels using CD31 antibody (scale bar = μm) and FACS quantification of CD31+ cells in dissociated tumours (n=3). **D)** Representative 3D reconstructions of mammary tumours imaged by multi-photon microscopy. Contrast in the tumour vasculature was achieved by tail-vein injection of blood quantum dots in the mice 20 minutes before imaging. Box plots represent the quantification of the vascular patency and vessel thickness of 4 replicates. * denotes $p<0.05$.

**Supplementary Figure 2. A)** Overview of the experimental design for the Drop-seq analysis on PyMT/WT and /ELF5 tumours including a summary of the main experimental characteristics (bottom table). **B)** Quality cut offs (dashed lines) applied for the single cell RNAseq analysis, percentage of mitochondrial genes (15%), number of Unique Molecular Identifiers (UMI) per cell (8,000) and number of genes identifies per cell (4,000). The resultant distribution after cut offs is shown in the insets**. C)** tSNE plot showing the distribution of cells per replicate. **D**) Feature and violin plots of Elf5 expression in the analysed PyMT/WT and PyMT/Elf5 tumours.

**Supplementary Figure 3. A)** Feature tSNE plots showing the expression of canonical markers of each of the main cell lineages, red epithelial, blue stroma and green immune. **B)** Visualisation (tSNE and dot plots) of the top differential genes for each of the defined clusters in the immune lineage (top) and in the stroma (bottom).

**Supplementary Figure 4. A)** Comparison of the principal component analysis before (left panel) and after (right panel) CCA alignment. **B)** tSNE plot of the K-means clustering of the CCA aligned data by CCA-cluster identity (left panel), and grouped by genotype (mid and right panels). **C)** Violin plots showing Elf5 and PyMT gene expression in each epithelial cell cluster. **D)** Dot plot of the expression levels of the top differential marker genes in each of the PyMT clusters coloured by genotype. The yellow rectangles highlight the top genes represented by each cluster. The size of the dots represents the percentage of cells/cluster that express each particular gene (pct. exp) and the colour gradient shows the level of expression for each gene/cluster. Note both colours are shown only when the cluster was populated similarly by both genotypes according to Fig. 2C.

**Supplementary Figure 5.** GSVA analysis for the H Hallmarks from MSigDB in the cancer epithelial clusters. Waterfall plots are presented for clusters mainly formed by one genotype and a heatmap is presented in the clusters formed by the two genotypes (according to Figure 2B). The variation score shows the hallmarks enriched (light blue, positive values) and decreased (dark blue, negative values) for each cluster.

**Supplementary Figure 6. A)** Cluster tree modelling the phylogenic relationship of the different clusters within the cancer-associated fibroblasts at different clustering resolutions. **B)** Gene-expression heatmap of the top expressed genes for each fibroblast lineage (FC). **C)** tSNE representation of the cell cycle stages of the cancer-associated fibroblasts as defined by gene expression signatures using tSNE coordinates. Circled area shows the cycling cluster characterised by a total absence of G1 cells. **D)** Violin plots displaying the classical marker genes that define CAFs for each of the three fibroblast clusters defined in resolution 0.1 (FC). **E**) Meta-signatures using the gene profiles from Bartoschek et al for cross-reference of the nomenclature previously used defining fibroblast subsets in the PyMT model: Vascular CAFs (vCAFs) corresponding with myofibroblasts with a contractile signature; matrix CAFs (mCAFs), described as the subpopulation with the strongest ECM signature, correspond with the secretory type of fibroblasts and in more extend to the ECM-CAFs within this group; cycling CAFs (cCAFs) completely overlap with the cell cycle signature, as shown in panel C and development CAFs (devCAFs) with an epithelial origin corresponds to a small population further clustered in resolution 1 (Figure 4E, cluster F5). **F)** Identification of a subset of fibroblasts that expressed the epithelial markers of PyMT and EpCAM, consistent with devCAFs in the Bartoschek et al publication.

**Supplementary Figure 7. A)** Waterfall plot of the GSVA analysis for the H Hallmarks from MSigDB in the CAF cluster defined by resolution 0.1 (CF); Cluster CF0 and CF1 were defined as “CAFs” and CF2 as “Myofibroblasts”. The variation score shows the hallmarks enriched (light blue, positive values) and not enriched (dark blue, negative values) for each cluster. **B)** tSNE representation of the CCA analysis of the CAF cell compartment revealing a similar polarisation between the two genotypes, PyMT/WT (blue) PyMT/ELF5 (red). **C)** Waterfall plot of the GSVA analysis for the H Hallmarks from MSigDB in the involution CAFs clusters defined by resolution 1 (clusters F2 and F3). The variation score shows the hallmarks enriched (light blue, positive values) and not enriched (dark blue, negative values) for each cluster. **D)** tSNE plot showing the assignment of each cell type for the analysis of cell to cell interactions using CellphoneDB. The circles correspond to the cells from the Elf5 genotype and the triangles are the cells coming from the WT genotype.

**Supplementary Figure 8. A)** SHG images of PyMT/WT and PyMT/ELF5 tumours assessed for differences in fibre orientation angle and quantification of frequency of fibre alignment ranging from the peak alignment. Different colours correspond to specific angles of orientation n=6 PyMT/WT and n=4 PyMT/ELF5. Inset shows the cumulative frequency of fibre alignment +/- 10 degrees from peak. **B)** Feature tSNE plots of all cell in tumours and metascore highlighting the expression of the invCAF markers (Col1a1, Mmp3 and Cxcl12) within the fibroblast compartment (red background, as annotated in Fig. 1). **C)** Low power (10X) magnification of representative images on the immunohistochemistry analysis of the COL1A1 (monomers), MMP3 and CXCL12 proteins on PyMT/WT and /ELF5 tumours. Black arrows denote cells with elongated structure and red arrows denote cells more infiltrated in the tumours and with a more rounded shape.

**Supplementary Figure 9.** Histological analysis of a mammary gland differentiation series featuring: virgin (V), late pregnancy (18 days *post-coitum*, 18DPC), established lactation (4 days *post-partum*, 4DPP), early involution (1 day involution, Inv. D1) and late involution (4 days involution, Inv. D4). First column shows representative images of Haemotoxylin and Eosin (H&E) staining of the mammary glands at low (10X) and high (20X) power magnification (inset). Representative images of the immunohistochemistry staining using COL1A1 (monomer), MMP3 and CXCL12 antibodies are shown at 20X magnification. Bottom panel shows the automatic FIJI-based quantification of the positive staining represented as % positively stained area.

**Supplementary Figure 10. Involution is a poor prognosis factor in the context of Elf5 expression in luminal breast cancer.**
**A)** Kaplan-Meier survival curves of breast cancer patients from the METABRIC cohort in relation to ELF5 and Involution signature expression. High and low classifications are based on top and bottom tertiles. ELF5 high patients (red) and Involution signature high patients (yellow); ELF5 low (green) and Involution signature low (blue) patients. Log-rank p values <0.05 are shown in red. Right panel shows the distribution of the PAM50 classified breast cancer subtypes in the top and bottom tertile of Elf5 or involution expressing patients. **B)** Cox multivariate analysis of METABRIC patient cohort to assess the hazard ratio of the involution signature independent of subtype, age and Ki67 status. **C)** Expression of the involution signature in patients categorised by high and low Elf5 expression in the METABRIC dataset (microarray units) and the TCGA dataset (RNAseq log10 units). Patients were segregated according to Elf5 expression levels based on tertiles, Elf5-high patients (red) were defined as the top-tertile and Elf5-low patients (green) as the bottom-tertile, **D)** Same analysis as panel A but only in Luminal patients. Right panels show the influence of the Involution signature in ELF5 low (top panel) and ELF5 high (bottom panel) luminal patients. Log-rank p values <0.05 are shown in red.

**Supplementary Video 1.** Video of intravital multiphoton microscopy of quantum dots (red) flowing through blood vessels in a tumour from 14-week-old PyMT/WT mouse

**Supplementary Video 2.** Video of intravital multiphoton microscopy of quantum dots (red) flowing through blood vessels in a tumour from 14-week-old PyMT/Elf5 mouse. Green cells correspond to GFP expression in the mammary tumour epithelium.

**Supplementary Table 1**
Functional annotation of each cell cluster from the tumour epithelium.

| Mammary lineage | Cluster number | HALLMARK<br>White bg: positive enrichment<br>Gray bg: negative enrichment | Function |
|---|---|---|---|
| Undifferentiated | 0 | Hedgehog Signalling | Cell proliferation |
| | | E2F Targets | |
| | | Mitotic Spindle | |
| | | G2M Checkpoint | |
| | | Reactive Oxygen Species Pathway | Inflammation/Immune Response |
| | | IL6 JAK STAT3 Signalling | |

| | | | |
|---|---|---|---|
| | | Allograft Rejection | |
| | | Interferon Gamma Response | |
| | | Interferon Alpha Response | |
| | | Inflammatory Response | |
| | | Xenobiotic Metabolism | |
| | | Angiogenesis | Interaction with TME |
| | | Notch Signalling | Cell differentiation |
| | | Epithelial Mesenchymal Transition | EMT |
| Luminal progenitors | 2 | Notch signalling | Cell Differentiation |
| | | TGF beta Signalling | |
| | | Mitotic Spindle | Cell Proliferation |
| | | G2M Checkpoint | |
| | | E2F Targets | |
| | | Myc Targets v1 | |
| | | Cholesterol Homeostasis | Cell Metabolism |
| | | Bile acid metabolism | |
| | | Peroxisome | |
| | 3 | Interferon Alpha Response | Immune response |
| | | Interferon Gamma Response | |
| | | Androgen Response | Hormone Response |
| | | Estrogen Response Late | |
| | | Estrogen Response Early | |
| | | IL2 STAT5 Signalling | |
| | | Hypoxia | Interaction with TME |
| | | Angiogenesis | |
| | | Coagulation | Inflammation |
| | | Inflammatory Response | |
| | | Reactive Oxygen Species Pathway | |
| | | Complement | |
| | | Epithelial Mesenchymal Transition | EMT |
| | | Mitotic Spindle | Cell Proliferation |
| | | G2M Checkpoint | |
| | | E2F Targets | |
| | | Myc Targets v1 | |
| | | Myc Targets v2 | |
| | | Hedgehog Signalling | |
| | | TGF beta Signalling | Cell Differentiation |
| | | Myogenesis | |
| | 4 | Cholesterol Homeostasis | Cell Metabolism |
| | | Fatty Acid metabolism | |
| | | Heme metabolism | |
| | | Epithelial Mesenchymal Transition | EMT |
| | | TNFA signalling via NFKB | Inflammation |
| | | IL6 JAK STAT3 Signalling | |
| | | Complement | |
| | | Coagulation | |

| | | | |
|---|---|---|---|
| Alveolar lineage | 1 | IL6 JAK STAT3 Signalling | Inflammation |
| | | Complement | |
| | | Interferon Alpha Response | |
| | | Estrogen Response Early | Hormone Response |
| | | Estrogen Response Late | |
| | | Epithelial Mesenchymal Transition | EMT |
| | | Mitotic Spindle | Cell proliferation |
| | | G2M Checkpoint | |
| | | E2F Targets | |
| | | Myc Targets v1 | |
| | | Myc Targets v2 | |
| | | Hedgehog Signalling | |
| | | Interferon Gamma Response | Immune Response |
| | 6 | IL6 JAK STAT3 Signalling | Inflammation |
| | | Complement | |
| | | Coagulation | |
| | | Cholesterol Homeostasis | Cell Metabolism |
| | | Hypoxia | Interaction with TME |
| | | Angiogenesis | |
| | | Reactive Oxygen Species Pathway | Oxidative stress |
| | | Estrogen Response Early | Hormone Response |
| | | Epithelial Mesenchymal Transition | EMT |
| | | Mitotic Spindle | Cell proliferation |
| | | G2M Checkpoint | |
| | | E2F Targets | |
| | | Myc Targets v1 | |
| | | Hedgehog Signalling | |
| | | Interferon Gamma Response | Immune Response |
| | | Interferon Alpha Response | |
| Hormone sensing lineage | 9 | Androgen Response | Hormone Response |
| | | Estrogen Response Late | |
| | | Estrogen Response Early | |
| | | IL2 STAT5 Signalling | |
| | | Hypoxia | Interaction with TME |
| | | Angiogenesis | |
| | | Coagulation | Inflammation |
| | | Complement | |
| | | Inflammatory Response | |
| | | IL6 JAK STAT3 Signalling | |
| | | Allograft Rejection | |
| | | Interferon Gamma Response | |
| Basal/Myoepithelial lineage | 7 | Complement | Inflammation |
| | | Coagulation | |
| | | Allograft Rejection | |
| | | Inflammatory Response | |
| | | Reactive Oxygen Species Pathway | |

| | | | |
|---|---|---|---|
| | | IL6 JAK STAT3 Signalling | |
| | | Interferon Gamma Response | |
| | | Angiogenesis | Interaction with TME |
| | | Hypoxia | |
| | | Myogenesis | Cell differentiation |
| | | Notch Signalling | |
| | | Epithelial Mesenchymal Transition | EMT |
| | 8 | Notch Signalling | Cell differentiation |
| | | TGF beta Signalling | |
| | | Myogenesis | |
| | | Epithelial Mesenchymal Transition | EMT |

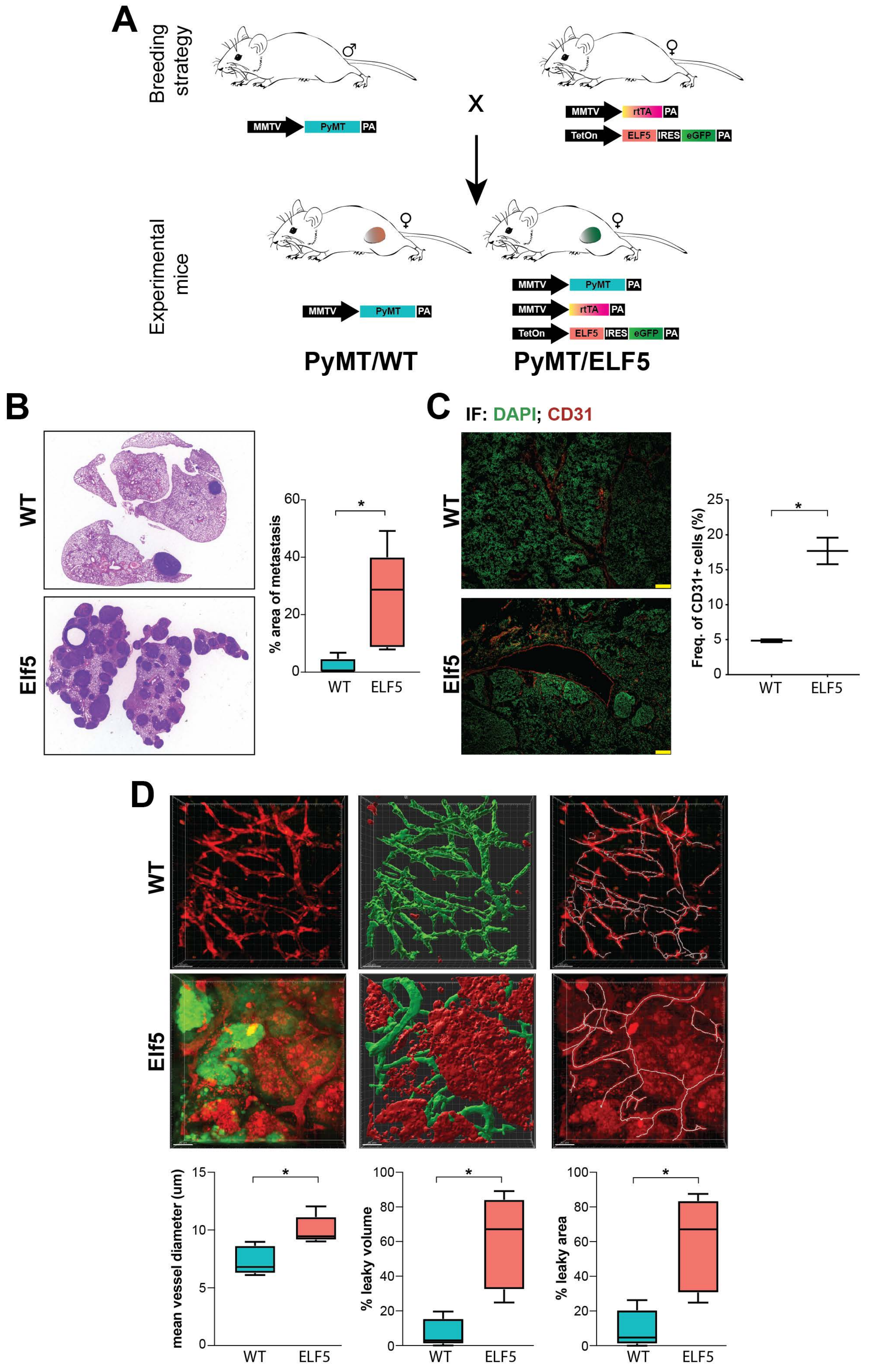

Supplementary Figure 1. Valdes-Mora et al. 2020

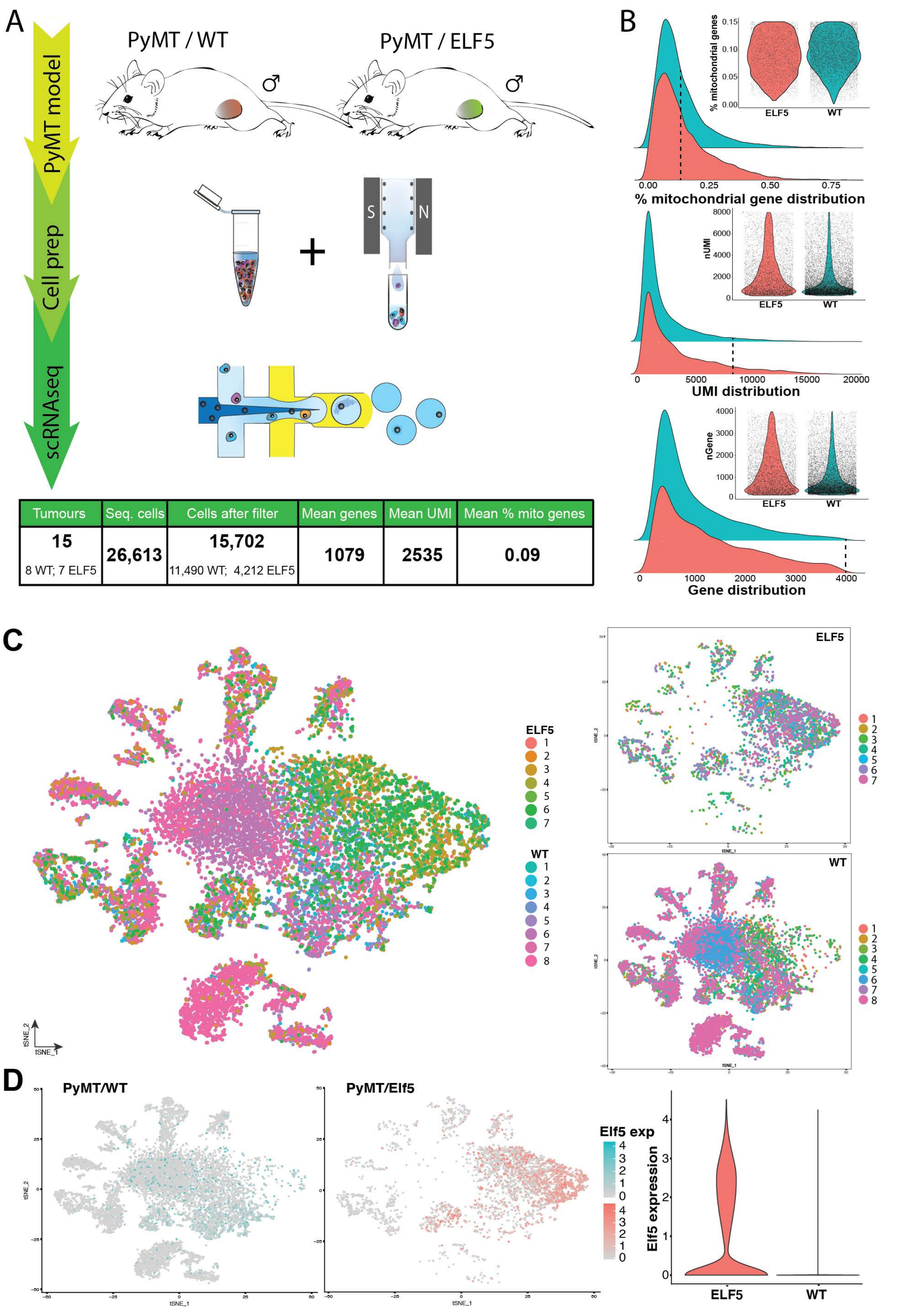

| Tumours | Seq. cells | Cells after filter | Mean genes | Mean UMI | Mean % mito genes |
|---|---|---|---|---|---|
| 15<br>8 WT; 7 ELF5 | 26,613 | 15,702<br>11,490 WT; 4,212 ELF5 | 1079 | 2535 | 0.09 |

Supplementary Figure 2. Valdes-Mora et al. 2020

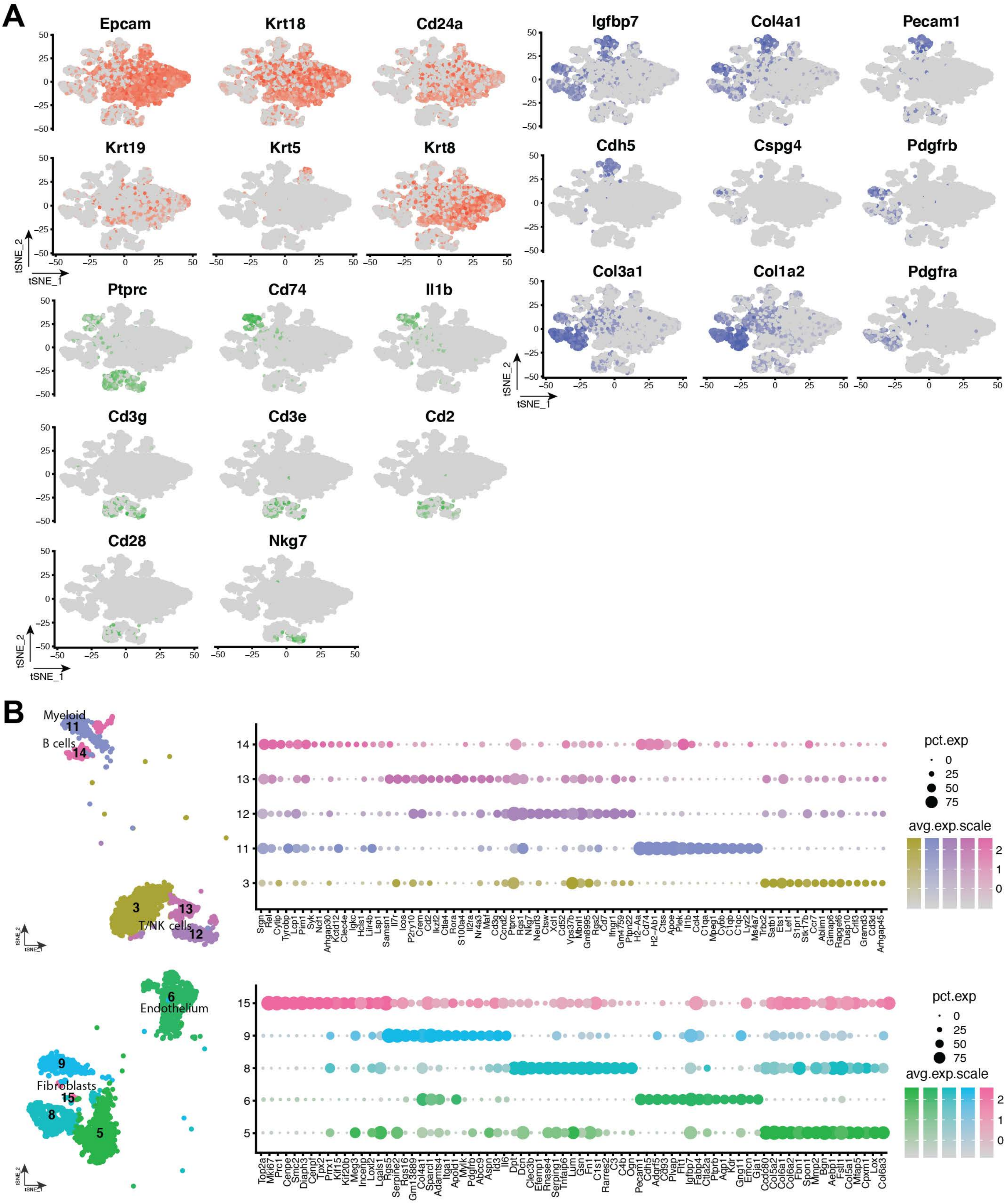

Supplementary Figure 3. Valdes-Mora et al. 2020

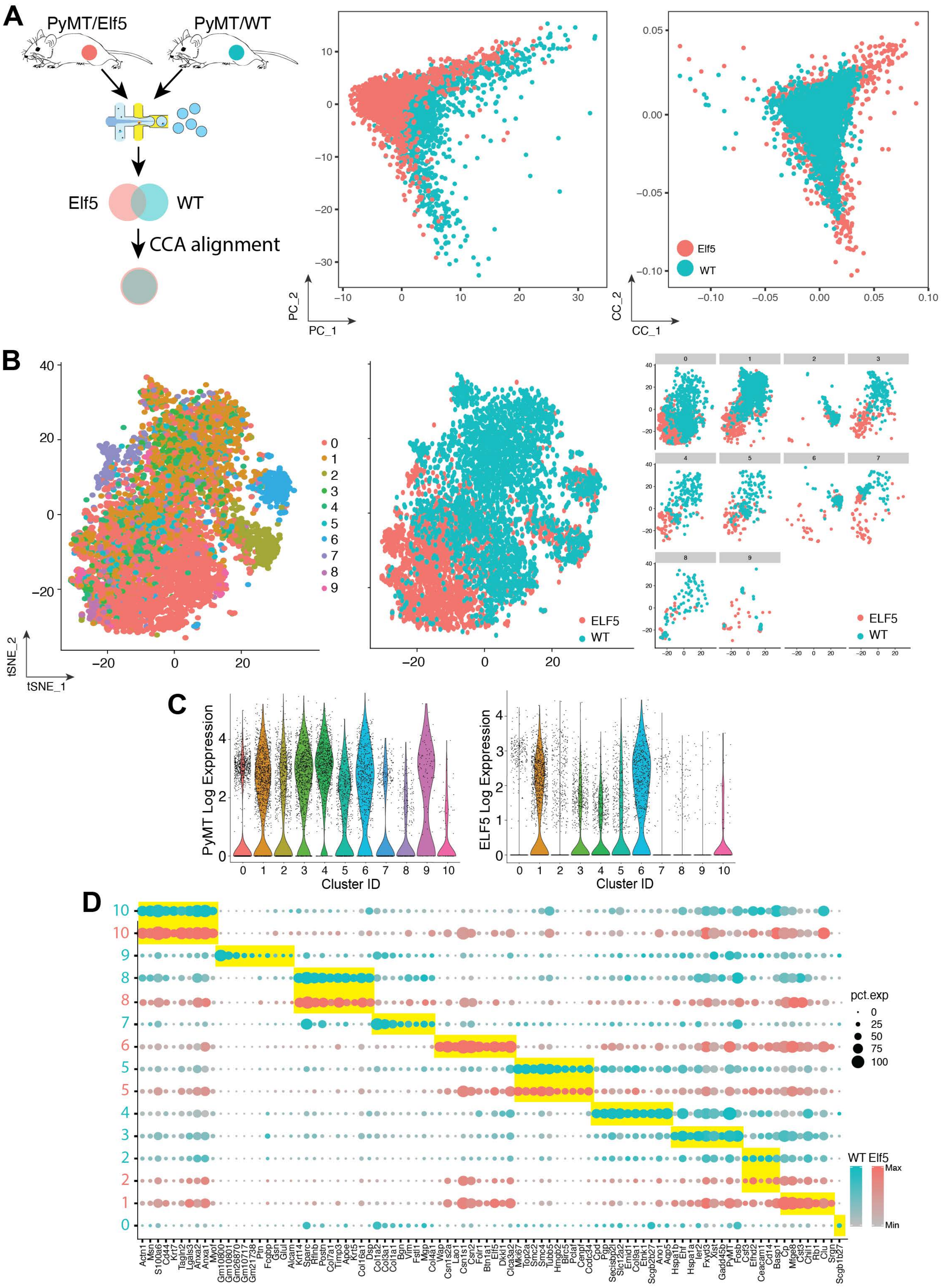

Supplementary Figure 4. Valdes-Mora et al. 2020

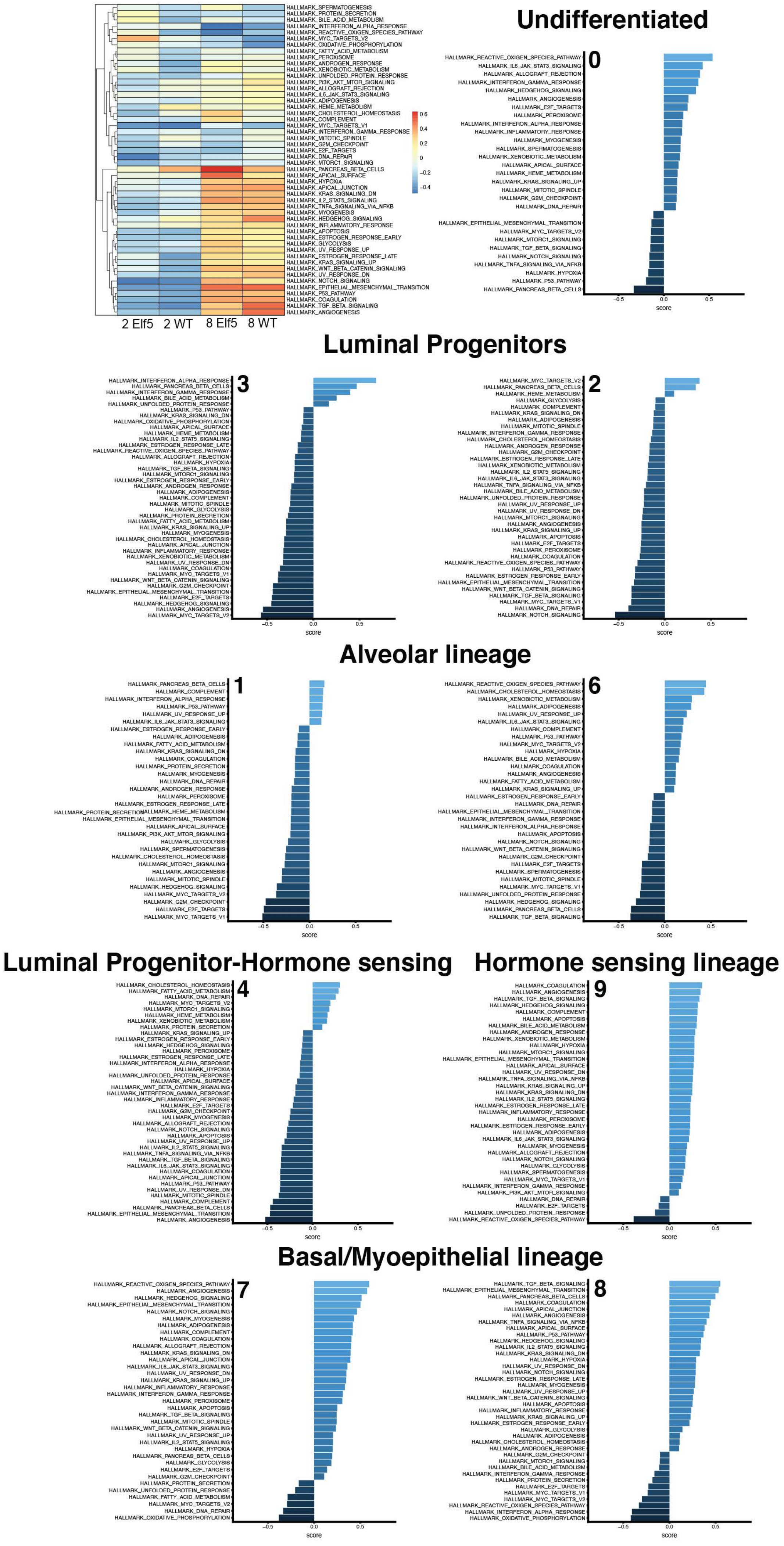

Supplementary Figure 5. Valdes-Mora et al. 2019

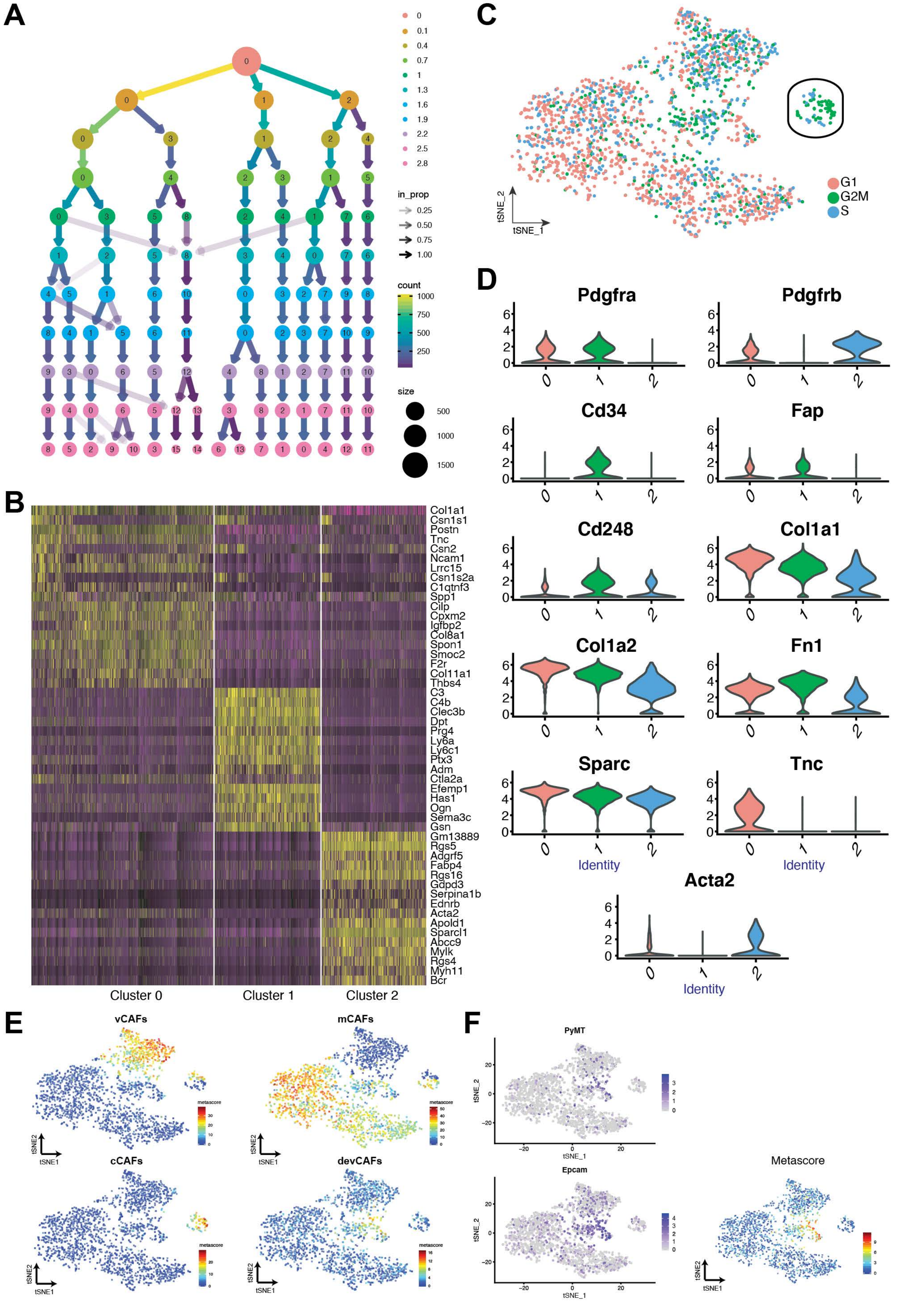

Supplementary Figure 6. Valdes-Mora et al. 2020

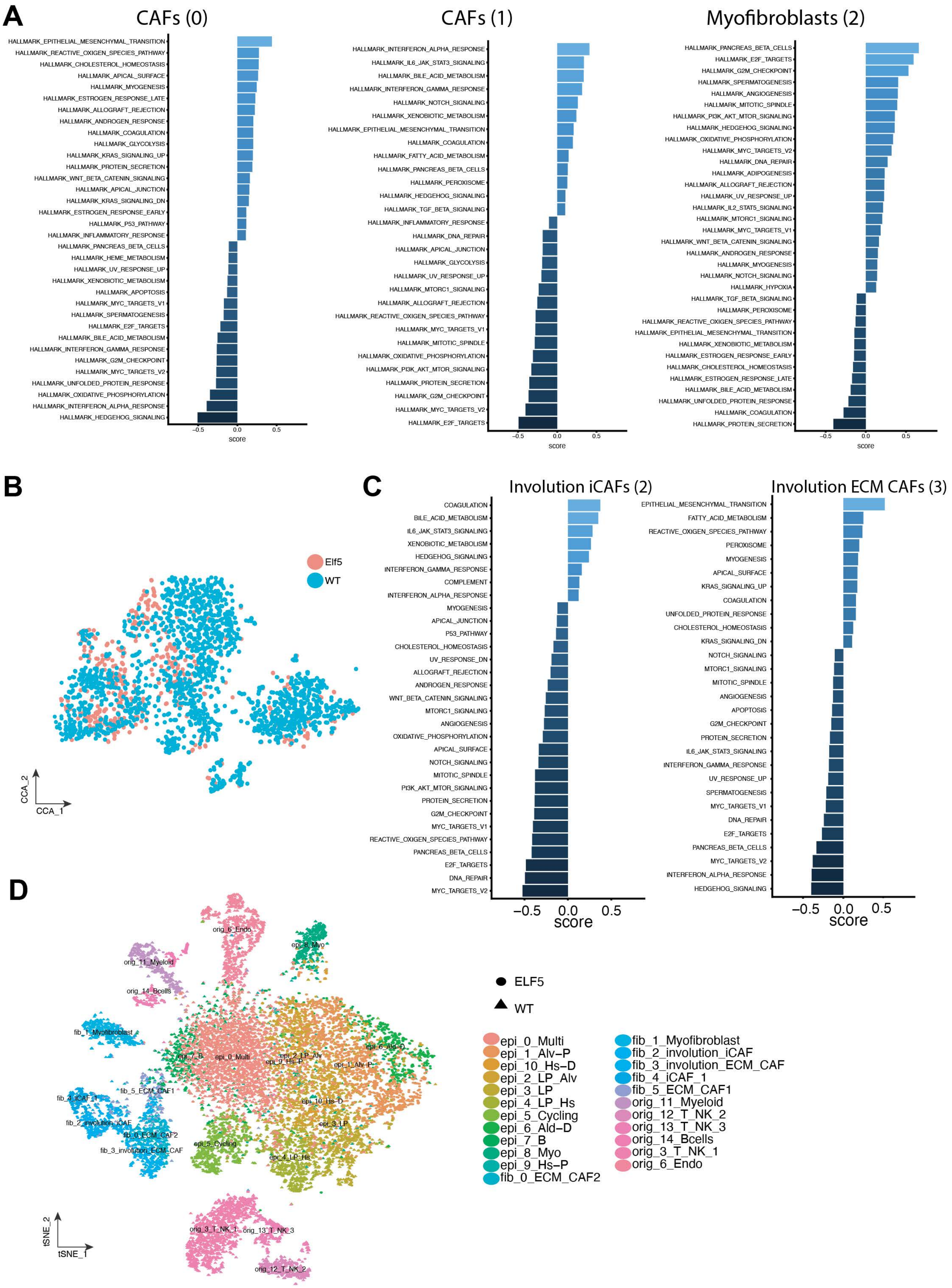

Supplementary Figure 7. Valdes-Mora et al. 2020

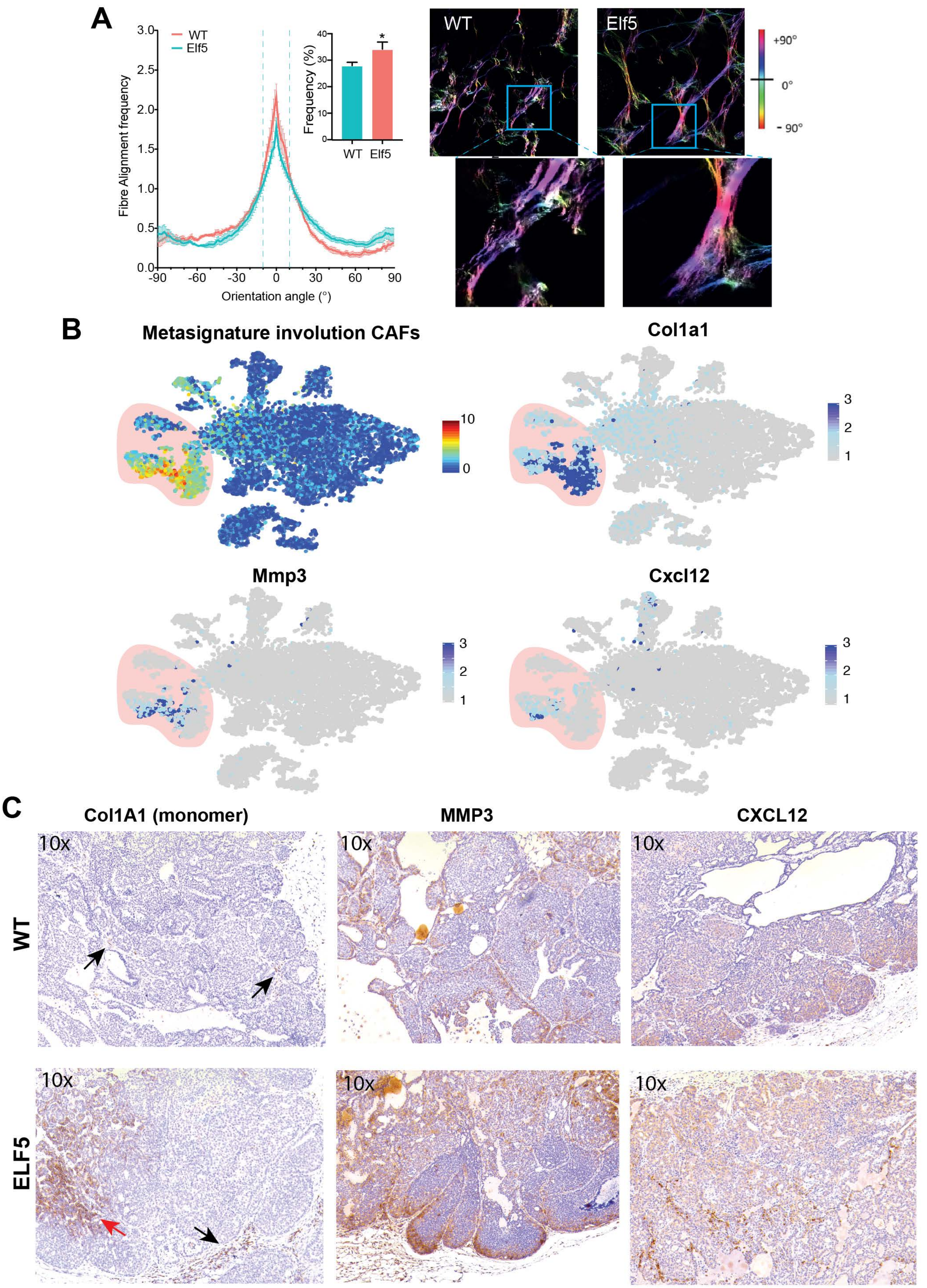

Supplementary Figure 8. Valdes-Mora et al. 2020

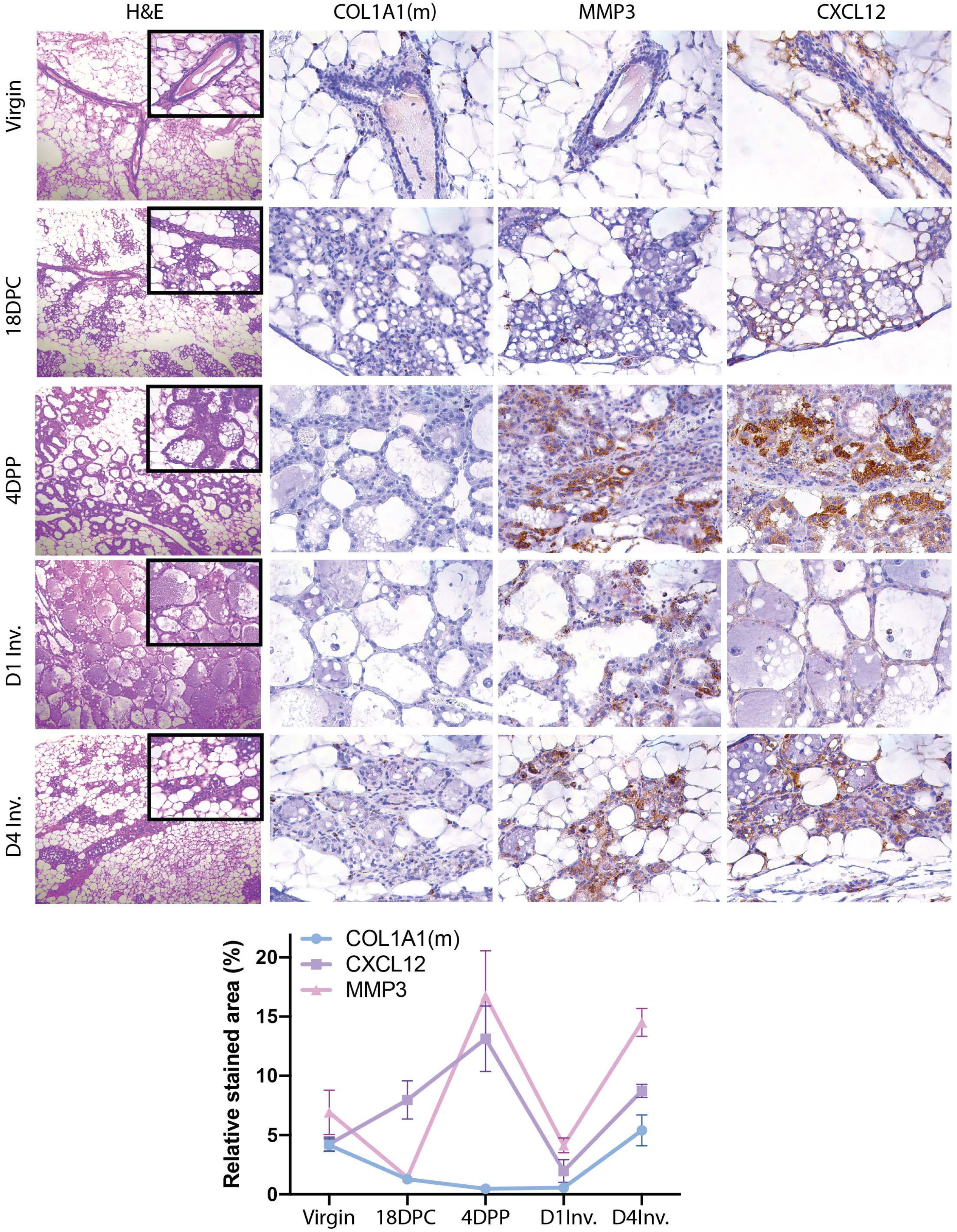

Supplementary Figure 9. Valdes-Mora et al. 2020

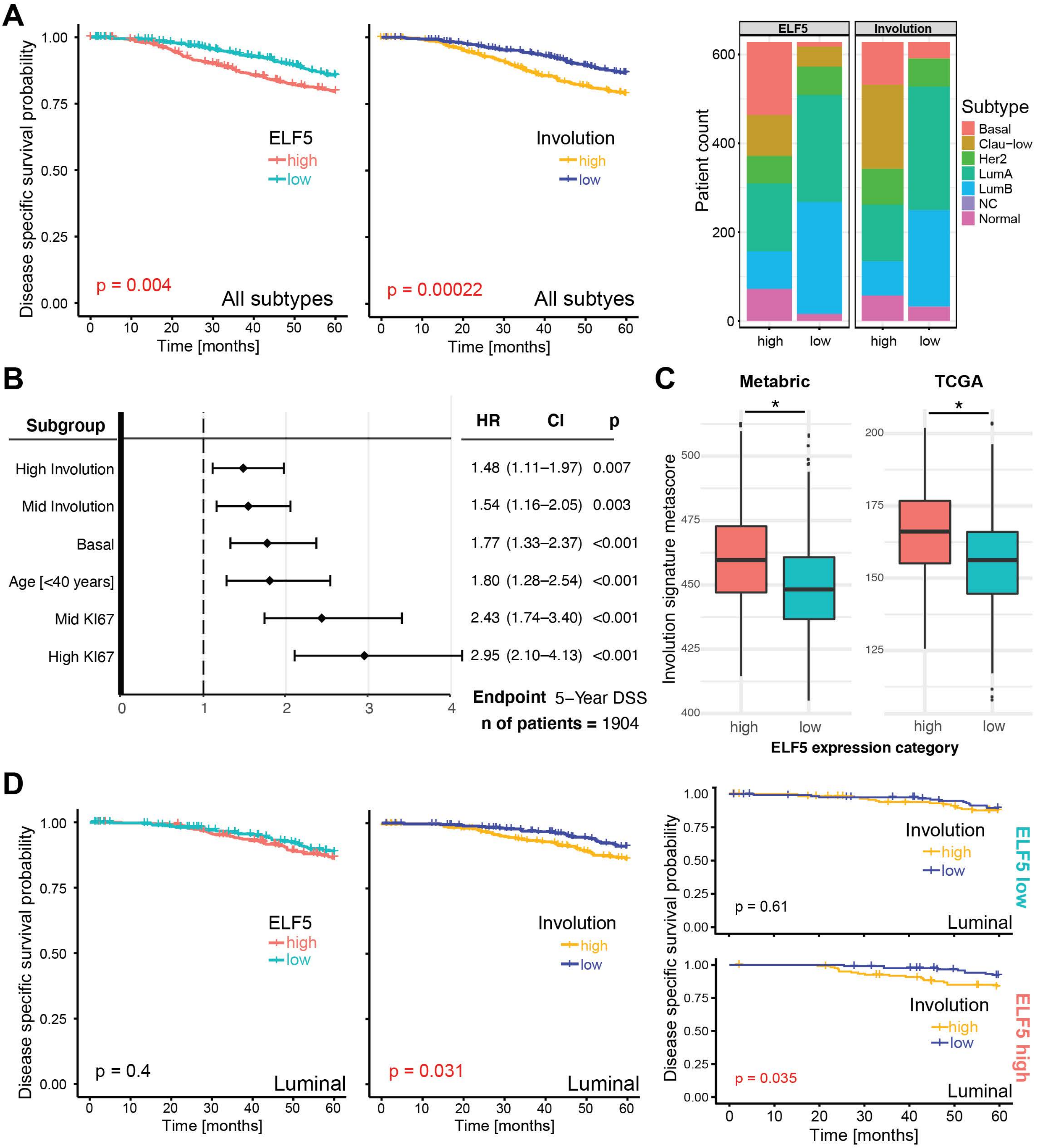

Supplementary Figure 10. Valdes-Mora et al. 2020
